# A nucleolar stress gene signature for quantitative scoring across multi-omics contexts

**DOI:** 10.64898/2026.04.12.718053

**Authors:** Jianxiong Chen, Shuai Xiao, Zhe Hao, Huimeng Xu, Xuemei Xu, Jun Zhou

**Author notes:** These authors contributed equally (Jianxiong Chen, Shuai Xiao).

## Abstract

The nucleolus is essential for ribosome biogenesis and cellular homeostasis, and its dysfunction can induce nucleolar stress, which has been implicated in cancer and other diseases. However, nucleolar stress is commonly inferred from morphological alterations or a limited set of functional assays, and quantitative approaches based on gene expression profiles remain lacking. Here, we integrate literature curation with multi-dataset screening to define a nucleolar stress gene signature and develop a nucleolar stress score (NuS) that is applicable across bulk transcriptomics, single-cell transcriptomics, proteomics, and spatial transcriptomics. Using this framework, we show in colorectal cancer models that oxaliplatin induces nucleolar stress, suppresses nascent rRNA synthesis, and activates p53 signalling, whereas these responses are attenuated in oxaliplatin-resistant cells. In combination with a ribosome biogenesis activity score (RiboSis), NuS captures related yet distinct dimensions of nucleolar function and stratifies tumors into functional states associated with distinct clinical outcomes. Furthermore, NuS-based analysis of perturbational transcriptomes enables prioritization of compounds with putative nucleolar stress-inducing activity. Collectively, this study establishes a quantitative framework for evaluating nucleolar stress and illustrates its applications in disease stratification and drug mechanism discovery.

## Introduction

The nucleolus is a membraneless nuclear compartment that serves as the center of ribosome biogenesis and a hub for multiple cellular processes ^1^. Its three-layered architecture, fibrillar center (FC), dense fibrillar component (DFC), and granular component (GC), supports the sequential steps of rRNA transcription, pre-rRNA processing, and ribosomal subunit assembly ^2^. Recent studies indicate that nucleolar organization is driven by liquid-liquid phase separation (LLPS) of proteins and nucleic acids, which underlies its structural plasticity and dynamic molecular exchange ^2,3^. Beyond ribosome production, the nucleolus also contributes to stress sensing, protein quality control, and genome organization ^4–7^. Perturbations in nucleolar structure or function that impair ribosome biogenesis are linked to a wide spectrum of human diseases, including ribosomopathies, cancer, and neurodegenerative disorders ^8,9^. These alterations are collectively termed nucleolar stress (also known as ribosomal stress), and robust quantitative approaches are needed to define its biological and pathological roles.

Nucleolar stress can be triggered by diverse stimuli, including metabolic perturbations, oxidative stress, aberrant gene expression, chemotherapeutic agents, and DNA damage, which ultimately converge on disruption of ribosome biogenesis (Figure 1a-b) ^9,10^. These insults activate multiple signaling pathways, most notably the canonical p53 pathway, leading to cell cycle arrest, apoptosis, autophagy, or senescence; however, p53-independent mechanisms are increasingly recognized (Figure 1c-d) ^9,11–14^. Currently, nucleolar stress is most often inferred from morphological alterations of nucleolar structure (Figure 1e) ^8^. In parallel, dysregulation of stress-associated signaling pathways, particularly activation of p53 signaling, is frequently used as molecular signatures ^15^, although its limited specificity, context dependence, and absence in many tumors constrain its utility as a universal marker. Importantly, impairment of nucleolar function, especially disruption of ribosome biogenesis, represents the most direct and mechanistically relevant evidence of nucleolar stress. However, such functional measurements are often not readily adaptable to rapid or large-scale evaluation.

**Figure 1.**
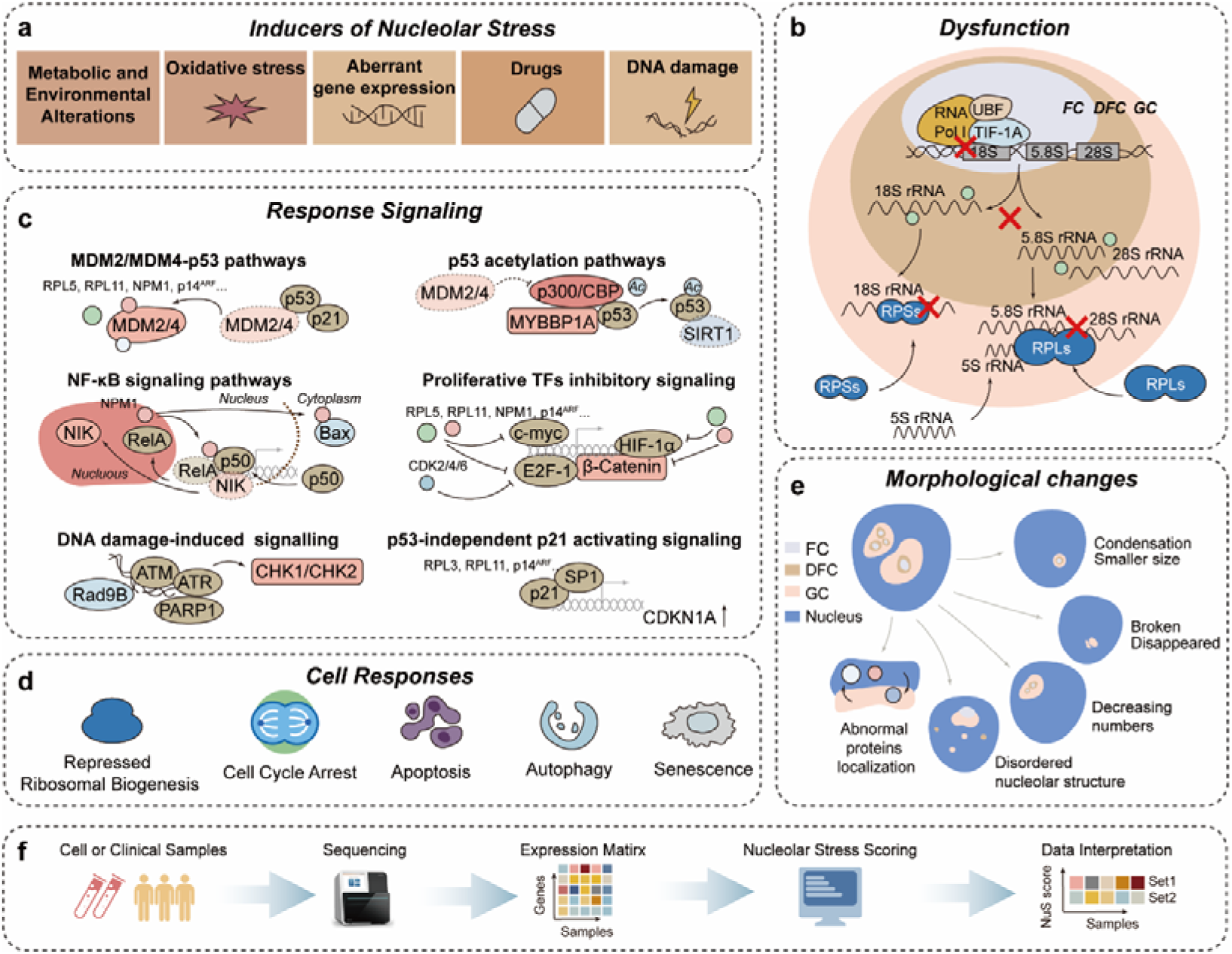
Nucleolar stress and the NuS scoring framework. (a) Major triggers of nucleolar stress, including metabolic/environmental perturbations, oxidative stress, dysregulated gene expression, pharmacological agents, and DNA damage. (b) Nucleolar organization and disruption under stress. The fibrillar center (FC), dense fibrillar component (DFC), and granular component (GC) support sequential steps of rRNA transcription, processing (18S, 5.8S, 28S), and ribosomal subunit assembly. Stress perturbs Pol I-dependent transcription (UBF, Pol I, TIF-IA), rRNA maturation, and ribosomal protein assembly, leading to nucleolar dysfunction. (c) Representative nucleolar stress signaling pathways, including MDM2/MDM4-p53, p53 acetylation (p300/CBP, MYBBP1A, SIRT1), NF-κB, repression of pro-proliferative transcriptional programs (c-Myc, HIF-1α, E2F1, β-catenin), DNA damage signaling (ATM/ATR-CHK1/CHK2), and p53-independent p21 (CDKN1A) activation. (d) Cellular outcomes, including reduced ribosome biogenesis, cell-cycle arrest, apoptosis, autophagy, and senescence. (e) Typical nucleolar morphological changes under stress, including condensation/shrinkage, fragmentation or loss, reduced nucleolar number, disorganization, and protein mislocalization. (f) NuS workflow: expression matrices from cell lines or clinical samples are used to compute NuS for quantitative interpretation of nucleolar stress states.

Recent efforts have attempted to quantify nucleolar states using morphology-based metrics, such as the nucleolar normality score^16^, which provide a useful framework for assessing nucleolar integrity. Nevertheless, not all forms of nucleolar stress are accompanied by overt structural alterations, and in some contexts morphological changes emerge only after the stress response has already been initiated, thereby limiting the sensitivity and general applicability of such approaches. Although gene sets have been developed to quantify ribosome biogenesis activity ^17,18^, no curated Gene Ontology (GO) term or KEGG pathway specifically captures the nucleolar stress response at the gene expression level. Moreover, whether suppression of ribosome biogenesis alone reliably reflects nucleolar stress severity remains unclear. Therefore, a curated gene signature and a robust scoring framework are needed to quantitatively evaluate nucleolar stress and to elucidate its functional relevance in disease (Figure 1f).

Gene set-based approaches enable quantitative inference of pathway activity from gene expression data by integrating coordinated expression patterns across predefined signatures ^19,20^. Such methods have been widely used to delineate cellular states, classify disease subtypes, and predict therapeutic responses. Here, we establish both full and core nucleolar stress-related gene sets through an integrative strategy combining literature curation with data-driven screening. Across bulk transcriptomic, proteomic, single-cell, and spatial transcriptomic datasets, we show that the resulting nucleolar stress score (NuS) provides a robust quantitative measure of nucleolar stress across diverse molecular contexts. We further demonstrate that ribosome biogenesis activity and nucleolar stress represent related yet distinct dimensions of nucleolar function, and that their combined assessment stratifies tumors into functionally distinct states with prognostic relevance. Finally, we show that NuS can serve as a useful framework for prioritizing anti-cancer compounds with nucleolar stress-inducing potential. Collectively, our work establishes a systematic approach for quantitative evaluation of nucleolar stress, providing a foundation for mechanistic studies and supporting translational applications across diverse disease contexts.

## Results

### Construction of a nucleolar stress-related gene signature

To address the lack of expression profile-based metrics for nucleolar stress, we combined literature curation, multi-dataset screening, and independent validation to derive a nucleolar stress gene signature (Figure 2a). We began with an extensive literature survey and compiled genes reported to respond to or regulate nucleolar stress. The initial curation yielded 131 candidate genes (47 upregulated and 84 downregulated), after excluding genes with inconsistent directionality across studies (Supplementary Tablès 1-2 and Supplementary Data 1). In addition, proteins reported to relocalize upon nucleolar stress (Supplementary Table 3), but lacking reproducible changes at the gene expression level, were not included.

**Figure 2.**
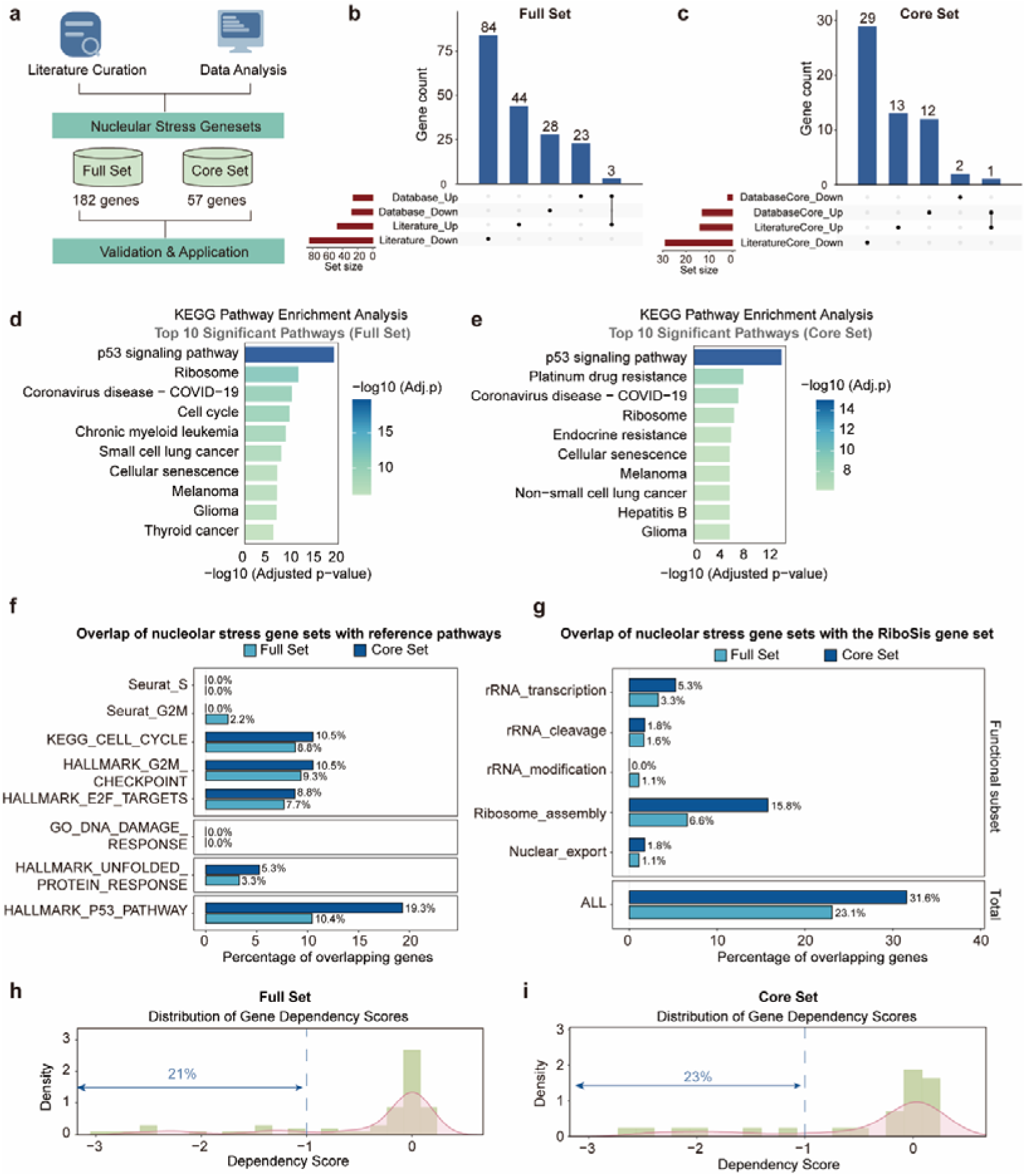
Construction and validation of a nucleolar stress gene signature. (a) Workflow for nucleolar stress signature construction. (b) Composition of the Full Set (182 genes). (c) Composition of the Core Set (57 genes). (d-e) KEGG pathway enrichment analysis of the Full Set (top 10 terms) and Core Set (top 10 terms). (f) Overlap analysis with cell cycle, DNA damage, endoplasmic reticulum (ER) stress, and p53 signaling gene sets. (g) Overlap with RiboSis genes. (h-i) DepMap dependency analysis. The percentage of upregulated genes meeting the essentiality threshold (dependency score < –1) is shown for the Full Set (h) and Core Set (i).

To expand the gene signature in a data-driven manner, we analyzed 17 transcriptomic datasets spanning multiple cell types treated with RNA polymerase I inhibitors, including CX-5461, BMH-21, and actinomycin D (Act D) (Supplementary Data 2). Differentially expressed genes (DEGs) were defined as |log_2_FC| > 1 with adjusted *P* < 0.05 (or *P* < 0.05 when adjusted values were unavailable) (Supplementary Data 2). For datasets containing multiple time points from the same cell line (GSE62963 and GSE198178), only DEGs with consistent directionality across time points were retained to improve robustness. In total, we identified 3,766 DEGs (1,277 upregulated and 2,489 downregulated; Supplementary Data 3). To prioritize genes most closely associated with nucleolar stress, we selected those recurrently altered in at least four independent datasets (Supplementary Figure 1), yielding 54 dataset-derived genes (26 upregulated and 28 downregulated; Supplementary Data 4).

We next integrated literature-derived and dataset-derived genes to construct a comprehensive nucleolar stress gene set, with genes showing discordant evidence between the two sources removed. This integration yielded a refined Full Set of 182 nucleolar stress-associated genes (70 upregulated and 112 downregulated; Figure 2b and Supplementary Data 5). To define a high-confidence Core Set, we applied more stringent criteria . Specifically, literature-derived genes were required to be supported by at least two independent studies, whereas dataset-derived genes were required to be differentially expressed in at least five independent datasets with consistent regulation under at least two distinct RNA polymerase I inhibitors. This filtering yielded 57 high-confidence genes (26 upregulated and 31 downregulated; Figure 2c and Supplementary Data 6).

Functional annotation of both gene sets by KEGG enrichment analysis highlighted pathways central to nucleolar stress biology, including p53 signaling, cell cycle regulation, and ribosome-related processes (Figure 2d-e). We next examined the extent to which nucleolar stress-associated genes overlapped with gene sets related to the cell cycle and other stress responses. Notably, fewer than 10% of nucleolar stress-associated genes overlapped with cell cycle genes, and the overlap with DNA damage and endoplasmic reticulum (ER) stress gene sets was even smaller (Figure 2f). These results argue against NuS simply reflecting generic cytotoxicity or broadly activated stress-response programs. By contrast, overlap with the p53 signaling pathway was more pronounced, reaching nearly 20%, suggesting a closer association between nucleolar stress and p53 signaling (Figure 2f). We further compared our gene sets with RiboSis genes, a previously defined metric reflecting ribosome biogenesis activity ^18^. Approximately 30% of genes overlapped, indicating that nucleolar stress is related to, but distinct from, impaired ribosome biogenesis (Figure 2g). Moreover, DepMap dependency analysis showed that approximately 20% of upregulated genes in both the Full Set and Core Set met the essentiality threshold (dependency score < −1) (Figure 2h-i), supporting a important role for nucleolar stress-associated programs in cellular function. Together, these Full and Core sets provide a curated framework for quantitative assessment of nucleolar stress across expression-based datasets.

### Establishing a high-confidence nucleolar stress score (NuS)

With the nucleolar stress gene signature established, we next developed a quantitative framework to estimate nucleolar stress across diverse expression datasets. Because analytical requirements differ across platforms, we first benchmarked and validated the scoring strategy in transcriptomic data before applying it to downstream analyses.

For bulk RNA-seq and microarray datasets, we evaluated single-sample gene set enrichment approaches, including GSVA and ssGSEA. ssGSEA is particularly suitable for single-sample or small-cohort comparisons, whereas GSVA is advantageous for large-scale screening. NuS was defined as the difference between enrichment scores for the upregulated and downregulated components of the signature (NuS = Score_up_ − Score_down_). As an initial test, we calculated NuS in A549 cells treated with the sapphyrin derivative PCI-2050 and the canonical nucleolar stress inducer Act D ^21^. Both GSVA and ssGSEA captured similar overall trends; however, ssGSEA showed clearer group separation and lower within-group variability, indicating greater stability in small-cohort analyses (Supplementary Figure 2). Act D treatment robustly increased NuS, supporting the ability of the score to capture nucleolar stress responses. Although the Core Set largely recapitulated the patterns observed with the Full Set, the Full Set showed greater overall robustness (Supplementary Figure 2). Therefore, unless otherwise noted, we used ssGSEA with the Full Set for small-cohort analyses.

We next performed internal validation using the 14 datasets compiled during signature screening (Supplementary Data 2). Across multiple cell types and compounds, NuS consistently distinguished nucleolar stress induced by different treatments and increased with treatment duration for the same compound (Supplementary Figure 3), demonstrating that the score captures both stimulus-specific and time-dependent stress responses.

We further validated NuS using independent published datasets representing mechanistically distinct nucleolar stress contexts. In U87MG cells, inhibition of IMPDH2 by mycophenolic acid (MPA) is known to trigger nucleolar stress and growth arrest ^22^. Consistent with this, MPA increased NuS, and this effect was attenuated by exogenous guanosine supplementation (>10 μM), in agreement with the original experimental findings ^22^ (Supplementary Figure 4a). In embryonic stem cells, depletion of the nuclear noncoding RNA LINE1 by antisense oligonucleotides (ASO-L1) has been shown to disrupt nucleolar organization ^23,24^; correspondingly, ASO-L1 increased NuS, whereas the sense control (SO-L1) did not ^24^ (Supplementary Figure 4b). Likewise, in liver samples from mice with inducible expression of the arginine-rich peptide (PR)_97_, a reported nucleolar stress inducer, NuS were elevated relative to non-induced controls ^25^ (Supplementary Figure 4c). Together, these results show that NuS provides a rapid and quantitative assessment of nucleolar stress across cellular contexts and perturbations. They further support the gene signature-based NuS as a sensitive and robust indicator of nucleolar stress.

### Oxaliplatin induces nucleolar stress in colorectal cancer (CRC)

We next focused on clinically relevant chemotherapeutics. Canonical rRNA transcription inhibitors such as CX-5461 and Act D are well-established nucleolar stress inducers with anti-tumor activity ^26,27^. Recent studies further reported that oxaliplatin and 5-fluorouracil (5-FU), first-line agents for CRC, can also trigger nucleolar stress ^28–36^.

To evaluate this, we first quantified NuS in CRC models. In HCT116 cells, 5-FU treatment increased NuS in a time-dependent manner (24-48 h; Supplementary Figure 5a). Oxaliplatin similarly elevated NuS in p53 wild-type HCT116 cells, but not in p53-mutant DLD1 cells (Supplementary Figure 5b), consistent with a p53-dependent nucleolar stress response to oxaliplatin. We next assessed nucleolar morphology in p53 wild-type HCT116 and HCT8 cells using nucleophosmin (NPM1; GC marker) and fibrillarin (FBL; DFC marker) ^3,37^. Super-resolution SIM imaging revealed that oxaliplatin markedly disrupted nucleolar architecture, characterized by nucleolar rounding, fragmentation, and nucleolar cap formation, whereas 5-FU induced minimal morphological changes under the same conditions (Figure 3a and Supplementary Figure 5c). Consistently, in a time-course analyses, oxaliplatin induced rapid nucleolar rounding as early as 4 h, resembling Act D and CX-5461, while 5-FU failed to produce overt structural changes within 24 h (Figure 3b).

**Figure 3.**
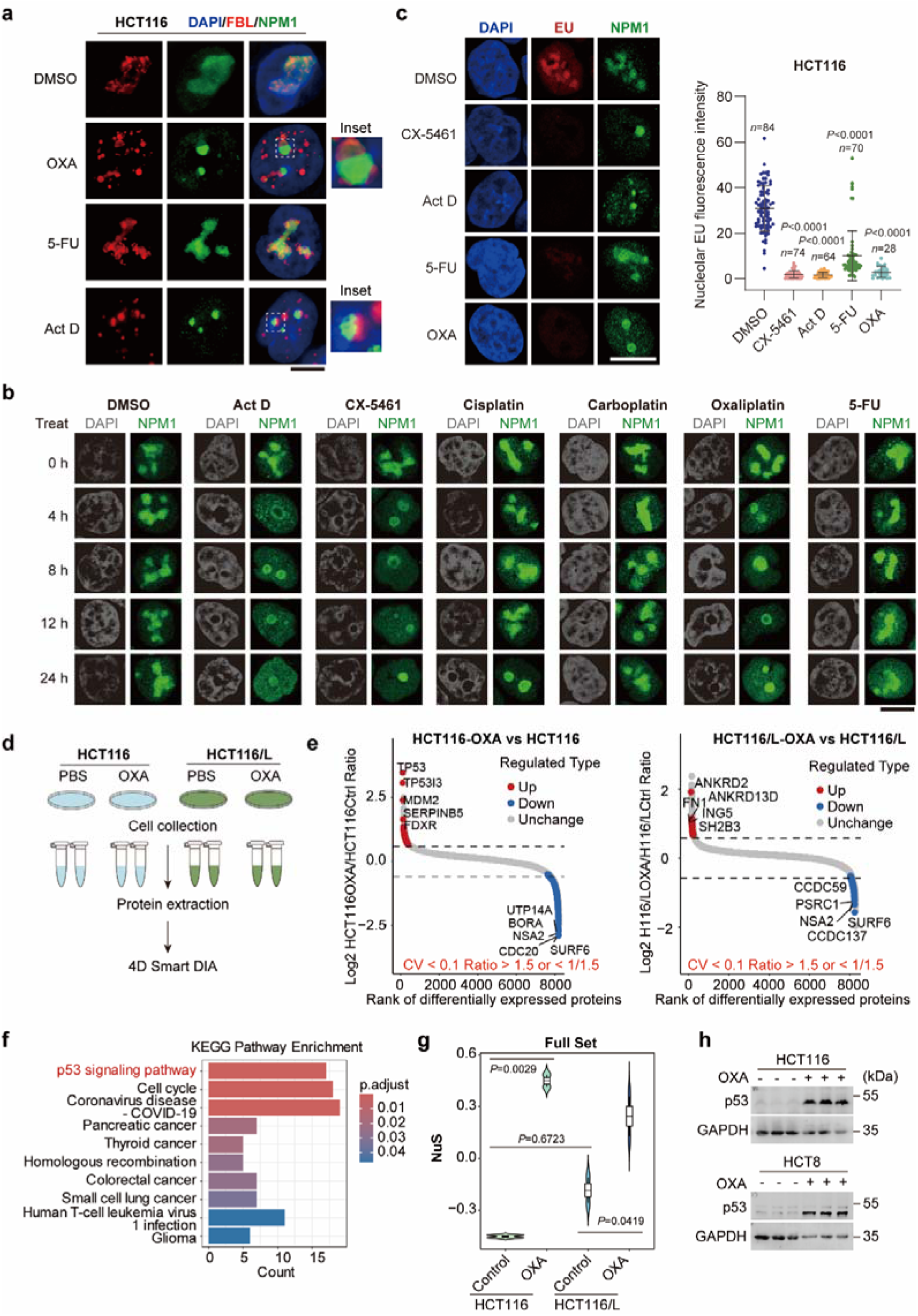
Oxaliplatin induces nucleolar stress in CRC cells. (a) HCT116 cells treated with DMSO, oxaliplatin (OXA, 12.5 μM) or 5-fluorouracil (5-FU, 1 μg/mL) for 24 h were stained for DAPI (blue), fibrillarin (FBL; red) and nucleophosmin (NPM1; green). Insets show enlarged nucleoli. Scale bar, 5 μm. (b) HCT116 cells treated with DMSO, Act D (50 ng/mL), CX-5461 (1 μM), cisplatin (16.7 μM), carboplatin (12.5 μM), OXA (12.5 μM) or 5-FU (1 μg/mL) for the indicated times were stained for DAPI (gray) and NPM1 (green). Scale bar, 5 μm. (c) 5-EU incorporation assay in HCT116 cells treated with the indicated agents. Representative images show DAPI (blue), EU (red) and NPM1 (green). Right, quantification of nucleolar EU fluorescence intensity; each dot represents one nucleolus, with *n* indicated. Kruskal–Wallis test. Scale bar, 5 μm. (d) Proteomic workflow for parental HCT116 and oxaliplatin-resistant HCT116/L cells treated with PBS or OXA (12.5 μM) for 24 h (*n* = 2 biologically independent samples per group) and analyzed by 4D Smart DIA mass spectrometry. (e) Volcano plots showing proteomic changes induced by OXA in HCT116 and HCT116/L cells. Proteins were classified using CV < 0.1 and fold change > 1.5 or < 1/1.5. (f) KEGG enrichment analysis of proteins differentially expressed in HCT116 cells after OXA treatment (FDR-adjusted *P* < 0.05). (g) NuS calculated from proteomic profiles of HCT116 and HCT116/L cells treated with PBS or OXA. One-way ANOVA followed by Tukey’s multiple-comparison test. (h) Immunoblot analysis of p53 in HCT116 and HCT8 cells treated with OXA for 24 h; GAPDH served as a loading control. Exact *P* values are indicated in the figure. Source data for c, g and h are provided in Source Data file.

Given that oxaliplatin has been proposed to induce nucleolar stress by inhibiting rRNA transcription ^31,33,35^, we next quantified 47S pre-rRNA levels. Oxaliplatin robustly reduced 47S pre-rRNA abundance, comparable to Act D and CX-5461, whereas 5-FU exerted a weaker effect in HCT116 and no detectable effect in HCT8 (Supplementary Figure 5d). Consistently, 5-ethynyl uridine (5-EU) incorporation assays further showed a pronounced reduction in nascent nucleolar RNA synthesis following oxaliplatin treatment, similar to Act D and CX-5461; in contrast, 5-FU caused only a modest decrease in HCT116 and had no detectable effect in HCT8 (Figure 3c and Supplementary Figure 5e). Notably, in HCT116 cells, 5-FU reduced rRNA transcription at 24 h without inducing overt nucleolar morphological changes, while NuS was increased, suggesting that NuS captures early transcriptional perturbations preceding structural remodeling.

To investigate whether oxaliplatin-induced nucleolar stress is linked to drug response, we generated oxaliplatin-resistant derivatives of HCT116 and HCT8 (Supplementary Figure 6a). Parental HCT116 cells and the resistant HCT116/L cells were treated with oxaliplatin for 24 h and analyzed by 4D label-free quantitative proteomics (Figure 3d). In parental cells, oxaliplatin increased p53 abundance and enriched p53-related pathways, whereas these responses were largely absent in HCT116/L cells (Figure 3e-f and Supplementary Figure 6b). Consistently, proteomics-derived NuS increased markedly in oxaliplatin-treated HCT116 cells, but only slightly in HCT116/L cells (Figure 3g and Supplementary Figure 6c), indicating an attenuated nucleolar stress response in the resistant state. Immunoblotting similarly confirmed p53 accumulation following oxaliplatin treatment in HCT116 and HCT8 cells (Figure 3h).

Finally, we also assessed ribosome biogenesis activity using the previously described RiboSis ^18^. Oxaliplatin suppressed ribosome biogenesis in both HCT116 and HCT116/L cells (Supplementary Figure 6d). Notably, baseline of ribosome biogenesis activity was lower in HCT116/L than in parental HCT116, whereas baseline NuS levels were comparable (Figure 3g and Supplementary Figure 6d), suggesting that reduced ribosome biogenesis may contribute to oxaliplatin resistance. Together, these results establish oxaliplatin as a potent inducer of nucleolar stress in CRC and demonstrate that NuS provides a robust and sensitive assessment across both transcriptomic and proteomic modalities.

### Differential ribosome biogenesis and nucleolar stress at single-cell resolution

Given the robust performance of NuS across bulk transcriptomic and proteomic datasets, we next evaluated its applicability at single-cell resolution. We analyzed 371,223 tumor and matched adjacent normal cells from 62 CRC patients ^38^. Unsupervised clustering resolved 88 cellular subsets spanning seven major lineages (Figure 4a). Using Seurat AddModuleScore, we quantified NuS and RiboSis at the single-cell levels and mapped their distributions across cell types (Figure 4b-c). Comparative analysis between tumor and normal compartments revealed coordinated but lineage-specific alterations in nucleolar function. In epithelial cells, NuS was significantly reduced in tumors, whereas mast, myeloid, plasma, stromal, and NKT/ILC populations exhibited increased NuS (Figure 4d). In contrast, RiboSis was increased in epithelial and stromal cells but decreased in the remaining cell types (Figure 4e). These patterns suggest increased biosynthetic demand in tumor-associated epithelial and stromal cells, while immune populations reside in a relatively stress-enriched state within the tumor microenvironment.

**Figure 4.**
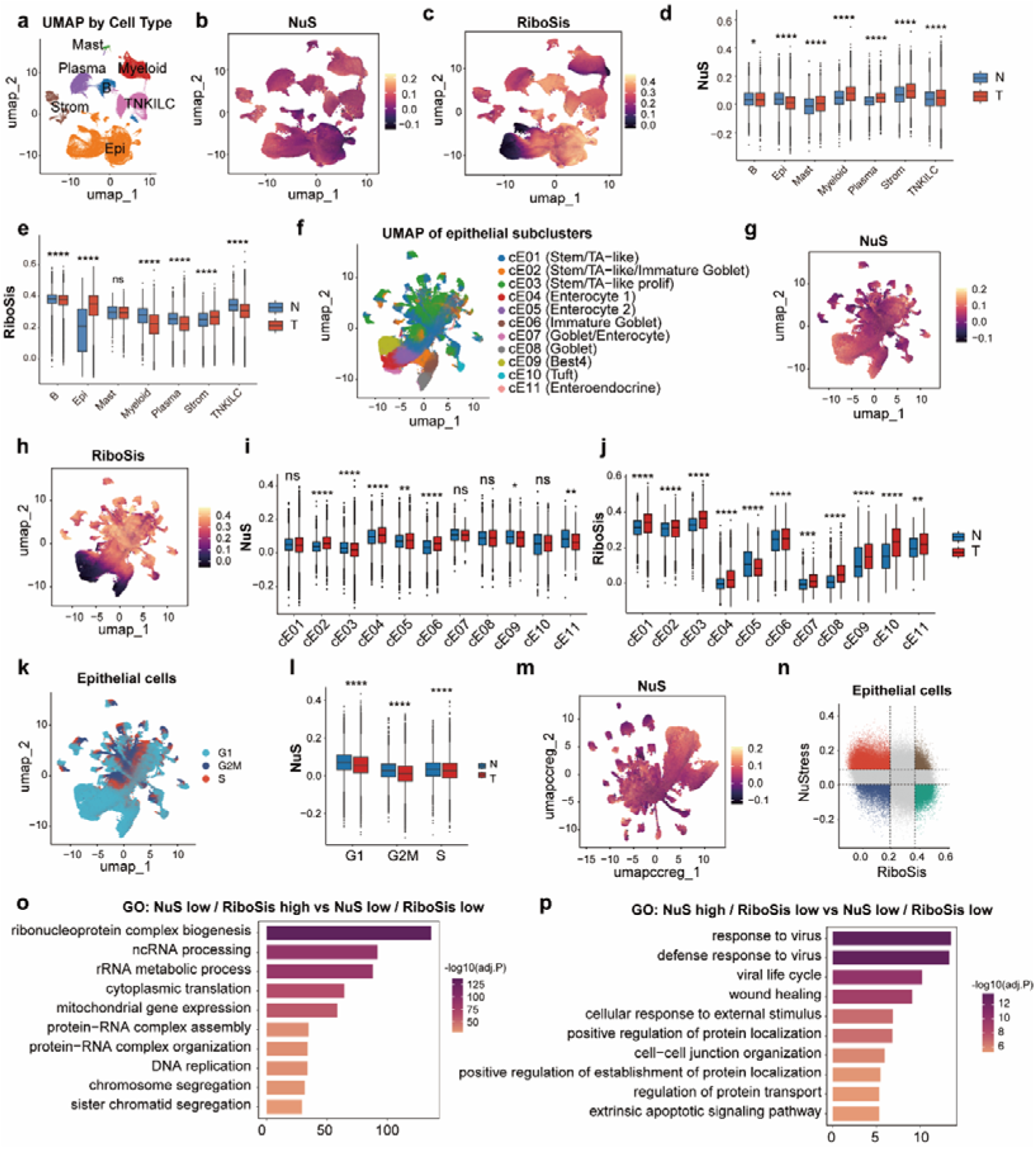
Single-cell transcriptomics reveals lineage-specific patterns of NuS and RiboSis in CRC. (a) UMAP of 371,223 single cells from 62 colorectal cancer (CRC) patients, annotated into seven major lineages—epithelial (Epi), stromal (Strom), T/NK/ILC, myeloid, plasma, B, and mast cells—based on the published classification. (b,c) UMAPs colored by NuS (b) and RiboSis (c). (d,e) Box plots comparing NuS (d) and RiboSis (e) between normal (N) and tumor (T) samples within each major lineage. Statistical significance was assessed using two-sided Wilcoxon rank-sum tests. (f) UMAP of epithelial cells showing 11 transcriptional subpopulations (cE01–cE11) according to the published annotation. (g,h) UMAPs showing the distribution of NuS (g) and RiboSis (h) across epithelial subpopulations. (i,j) Bar plots showing mean ± s.e.m. of NuS (i) and RiboSis (j) across epithelial subpopulations in normal (N) and tumor (T) samples. Statistical significance between N and T within each subpopulation was assessed using two-sided Wilcoxon rank-sum tests. (k) UMAP of epithelial cells colored by cell-cycle phase (G1, S, and G2/M), as determined by Seurat CellCycleScoring. (l) Box plots comparing NuS between normal and tumor epithelial cells within each cell-cycle phase. Statistical significance was assessed using two-sided Wilcoxon rank-sum tests. (m) UMAP of epithelial cells colored by NuS after regression of cell-cycle effects (S.Score and G2M.Score). (n) Quadrant plot stratifying epithelial cells into four groups according to NuS and RiboSis levels. Dashed lines indicate the thresholds used for stratification. (o) Gene Ontology (GO) enrichment analysis of genes upregulated in NuS-low/RiboSis-high epithelial cells relative to NuS-low/RiboSis-low cells. (p) GO enrichment analysis of genes upregulated in NuS-high/RiboSis-low epithelial cells relative to NuS-low/RiboSis-low cells. \**P* < 0.05, \*\**P* < 0.01, \*\*\**P* < 0.001, \*\*\*\**P* < 0.0001.

Given the pronounced tumor–normal differences in epithelial cells, we next examined epithelial heterogeneity in greater detail. Epithelial cells were further resolved into 11 transcriptional programs based on the published annotation ^38^ (Figure 4f). Both NuS and RiboSis varied substantially across epithelial subtypes, with RiboSis showing greater variability (Figure 4g-h and Supplementary 7a-d). Notably, ribosome biogenesis was elevated in nearly all tumor-associated epithelial programs except Enterocyte 2, whereas NuS exhibited heterogeneous and subtype-specific patterns (Figure 4i-j and Supplementary Figure 7a-d), suggesting that nucleolar stress is not a simple linear consequence of inhibited ribosome biogenesis.

As cell-cycle state may influence NuS and differ across epithelial subpopulations, we next examined whether the observed NuS patterns were primarily attributable to cell-cycle heterogeneity. We first analyzed cell-cycle phase distributions in epithelial cells (Figure 4k). NuS showed a modest negative correlation with both S-phase and G2/M scores (Supplementary Figure 7e,f). Consistent with this, NuS was lower in cells assigned to S and G2/M phases than in G1-phase cells (Supplementary Figure 7g), indicating that cell-cycle progression is associated with variation in NuS. However, when epithelial cells were stratified by cell-cycle phase, tumor-derived epithelial cells still exhibited lower NuS than their normal counterparts within each phase (Figure 4l). Moreover, regression of S.Score and G2M.Score did not abolish NuS variation across epithelial cells (Figure 4m and Supplementary Figure 7h). Together, these findings indicate that although NuS is influenced by cell-cycle state, the tumor-associated differences in NuS cannot be explained solely by cell-cycle effects.

To further dissect the sources of NuS and RiboSis heterogeneity, we stratified epithelial cells into distinct quadrants according to NuS and RiboSis levels (Figure 4n) and performed differential expression and pathway enrichment analyses. High-RiboSis epithelial cells were preferentially enriched for ribosome biogenesis-related programs (Figure 4o). In contrast, high-NuS cells were enriched for gene programs associated with responses to viral infection, external stimuli, and apoptotic signaling (Figure 4p). Furthermore, comparisons across normal and tumor epithelial cells, as well as across epithelial subpopulations, revealed clearly distinct distribution patterns for NuS and RiboSis (Supplementary Figure 7i-j). These findings suggest that NuS and RiboSis capture separable dimensions of nucleolar function, rather than simply representing opposing biological states. Collectively, these results demonstrate that NuS provides a robust single-cell assessment of nucleolar stress responses and captures functional heterogeneity beyond that explained by ribosome biogenesis alone, thereby complementing RiboSis.

### Spatial transcriptomics reveals heterogeneous nucleolar function

Although bulk and single-cell transcriptomics capture ribosome biogenesis and nucleolar stress programs, they lack spatial context. We therefore applied our framework to spatial transcriptomic profiles from four patients with CRC, each with paired primary tumors and liver metastases ^39^ (Figure 5a). Following unsupervised clustering, spatial regions were annotated according to the corresponding H&E images, showing strong concordance between histological features and transcriptional patterns (Figure 5b).

**Figure 5.**
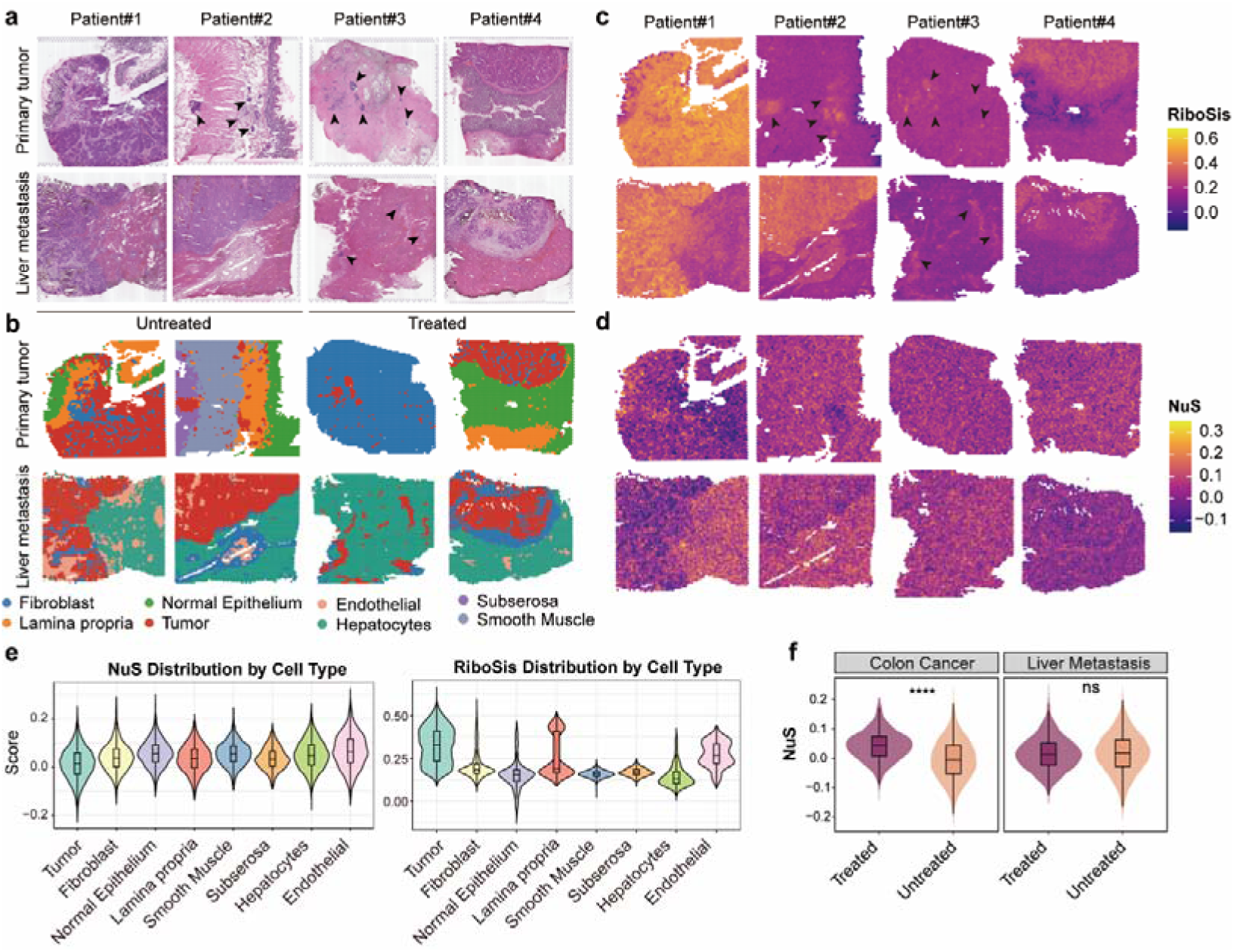
Spatial transcriptomics reveals heterogeneous nucleolar function in primary and metastatic CRC. (a) H&E-stained sections from four CRC patients with paired primary tumors and liver metastases. Arrows indicate tumor regions (top, primary; bottom, liver metastasis). (b) Spatial maps of annotated tissue regions guided by H&E morphology and clustering, including fibroblast, lamina propria, tumor, normal epithelium, hepatocytes, endothelial cells, subserosa, and smooth muscle. Patients #1-2 are untreated (left) and patients #3-4 received neoadjuvant therapy (right). (c-d) Spatial distribution of RiboSis (c) and NuS (d) across tissue sections. Warmer colors indicate higher scores. (e) Violin plots with overlaid box plots showing RiboSis (left) and NuS (right) across annotated tissue regions. (f) Violin plots with overlaid box plots showing NuS in tumor regions stratified by treatment status (untreated vs neoadjuvant-treated) in primary tumors (left) and liver metastases (right). Statistical significance was assessed using two-sided Wilcoxon rank-sum tests. ns, not significant; \*\*\*\**P* < 0.0001.

We next caculated RiboSis and NuS for each spatial spot using Seurat AddModuleScore and visualized their distributions across sections. Both scores exhibited substantial spatial heterogeneity (Figure 5c-d). Notably, ribosome biogenesis activity was strongly enriched in tumor regions and showed high spatial specificity, including within small tumor microlesions (Figure 5c). By contrast, NuS varied across tissue compartments and was less restricted to tumor regions (Figure 5d). Consistent with the single-cell transcriptomics analyses, RiboSis was higher in tumor than in non-tumor regions, while the NuS was lower in tumors (Figure 5e). The close spatial correspondence between high RiboSis and tumor cell-rich regions suggests that RiboSis may aid spatial delineation of CRC, with potential value for defining tumor boundaries and guiding sampling strategies.

We next assessed the impact of neoadjuvant XELOX therapy (capecitabine plus oxaliplatin). Treatment markedly increased NuS in primary tumors (Figure 5f), while reducing RiboSis in both primary tumors and liver metastases (Supplementary Figure 8a). Before treatment, NuS was lower in primary tumors than in metastatic lesions; however, after neoadjuvant therapy, NuS in primary tumors increased to levels exceeding those in metastases (Figure 5f and Supplementary Figure 8b). In contrast, liver metastases consistently exhibited lower RiboSis than primary tumors, regardless of treatment status (Supplementary Figure 8b). Further pathway analysis showed that metastatic tumors were characterized by enhanced metabolic remodeling and downregulation of ribosome biogenesis-related pathways (Supplementary Figure 8c), suggesting that metastatic lesions may adopt a less proliferative but more adaptive state within the hepatic microenvironment. Together, these results support the applicability of NuS and RiboSis to spatial transcriptomics and show that these complementary metrics capture both the spatial organization of tumor-associated niches and dynamic changes associated with therapy and metastatic progression.

### NuS and RiboSis define closely related but distinct aspects of nucleolar function in cancer

To systematically characterize nucleolar function across human cancers, we computed NuS and RiboSis using ssGSEA in treatment-naïve TCGA RNA-seq cohorts together with normal tissues from GTEx. To minimize treatment-related confounding, samples annotated as having received neoadjuvant therapy were excluded (Supplementary Figure 9). At the pan-cancer level, NuS was consistently lower in tumors than in normal tissues (Figure 6a), in agreement with our single-cell observations. In contrast, RiboSis was broadly elevated in tumors (Figure 6a), consistent with increased ribosome biogenesis in cancer ^18^.

**Figure 6.**
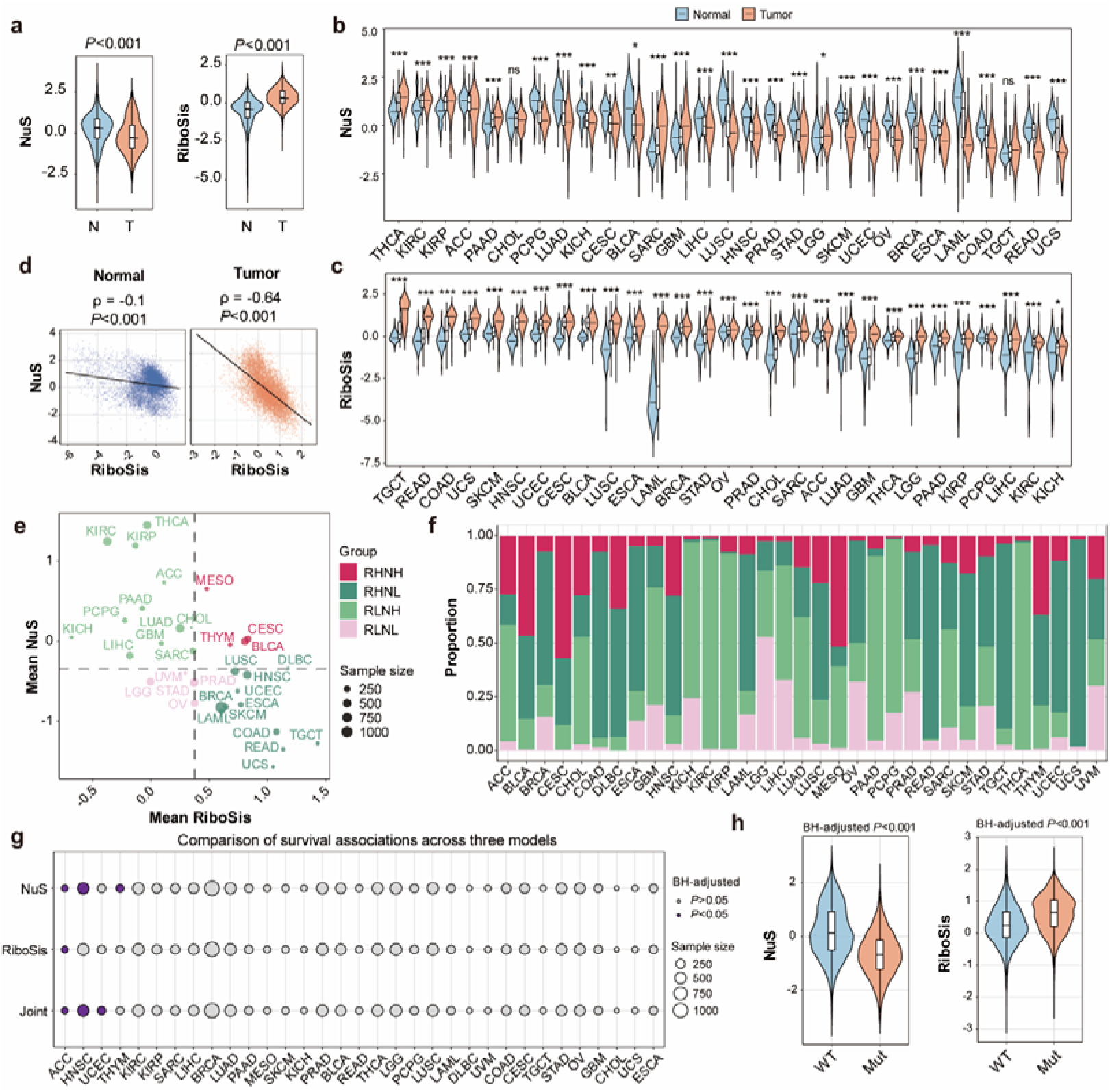
NuS and RiboSis captures nucleolar functional states and heterogeneity across cancers. (a) Violin plots comparing NuS (left) and RiboSis (right) between normal (N) and tumor (T) samples across TCGA pan-cancer cohorts and GTEx. Statistical significance was assessed using a two-sided Wilcoxon rank-sum test. (b, c) Pan-cancer distribution of NuS (b) and RiboSis (c) across individual cancer types, comparing normal (blue) and tumor (orange) samples. For each cancer type, differences between groups were evaluated using a two-sided Wilcoxon rank-sum test (wilcox.test). Only comparisons with sufficient sample size (minimum n ≥ 3 per group) were tested. (d) Correlation between NuS and RiboSis in normal and tumor tissues. Each dot represents one sample. Spearman correlation coefficients (ρ) and corresponding P values are shown. (e, f) Pan-cancer landscape of nucleolar functional states defined by NuS and RiboSis. In (e), each dot represents one cancer type, positioned by the mean NuS and mean RiboSis values across tumor samples; dot size is proportional to sample number. In (f), tumor samples within each cancer type were classified into four groups using median cutoffs of NuS and RiboSis: RiboSis-high/NuS-high (RHNH), RiboSis-high/NuS-low (RHNL), RiboSis-low/NuS-high (RLNH), and RiboSis-low/NuS-low (RLNL). Stacked bars show the proportion of each subtype across cancer types. (g) Prognostic relevance of NuS, RiboSis, and their joint stratification across cancer types. Dot plot shows the significance of overall survival associations for each model; dot size indicates sample number, and the dashed line marks P = 0.05. Survival differences were assessed by log-rank test, and multiple-testing correction was performed using the Benjamini-Hochberg method. (h) Comparison of NuS (left) and RiboSis (right) between TP53 wild-type (WT) and TP53-mutant (Mut) tumors. Statistical significance was assessed using a two-sided Wilcoxon rank-sum test. ns, not significant; * *P* < 0.05; ** *P* < 0.01; *** *P* < 0.001.

Across individual cancer types, ribosome biogenesis was broadly increased, whereas NuS exhibited substantially greater heterogeneity (Figure 6b-c). Several malignancies, including thyroid carcinoma, kidney renal clear cell carcinoma, kidney renal papillary cell carcinoma, and adrenocortical carcinoma, displayed relatively high NuS levels (Figure 6b). Despite partial coupling, NuS and RiboSis showed only moderate correlation, with stronger associations observed in tumor tissues than in normal tissues (Figure 6d). This indicates that nucleolar stress and ribosome biogenesis are partially uncoupled, reflecting distinct yet overlapping regulatory programs. Accordingly, NuS captures biological dimensions beyond ribosome biogenesis alone.

To further delineate nucleolar functional states, we mapped cancers within a two-dimensional space defined by NuS and RiboSis. This revealed diverse nucleolar phenotypes, ranging from tumors with high ribosome biogenesis (e.g., TGCT, COAD/READ, and UCS) to those with elevated nucleolar stress (e.g., THCA, KIRC, KIRP; Figure 6b-c). We then stratified tumors into four nucleolar functional subtypes using median cutoffs of both metrics: RiboSis-high/NuS-high (RHNH), RiboSis-high/NuS-low (RHNL), RiboSis-low/NuS-high (RLNH), and RiboSis-low/NuS-low (RLNL) (Figure 6e). This classification captured both differences between tumor types and substantial heterogeneity within individual cancer types (Figure 6f), highlighting the complexity of nucleolar functional states across patients. We next evaluated the clinical relevance of these metrics. NuS alone was significantly associated with patient prognosis in multiple cancers, including ACC, HNSC, and THYM, whereas RiboSis showed prognostic value in ACC. Importantly, the combined nucleolar functional subtypes provided improved prognostic stratification in ACC, HNSC, and UCEC (Figure 6g and Supplementary Figure 10), underscoring the added value of integrating both axes.

Given the central role of TP53 in nucleolar stress signaling, we further examined its relationship with these metrics. Tumors harboring TP53 mutations exhibited significantly lower NuS and higher RiboSis scores (Figure 6h and Supplementary Figure 11a–b), consistent with impaired nucleolar stress responses and enhanced ribosomal biogenesis activity. In CRC, NuS and RiboSis were also associated with Consensus Molecular Subtypes (CMS)^40^: NuS was lowest in CMS2 tumors, whereas RiboSis was highest in this subtype (Supplementary Figure 11c,d), in line with the known enrichment of ribosome biogenesis and MYC signaling in CMS2.

To determine whether NuS merely reflects canonical TP53 signaling, we performed TP53-stratified analyses across TCGA tumors. Notably, the inverse association between NuS and RiboSis was preserved in both TP53 wild-type and TP53-mutant tumors, and remained particularly strong in the TP53-mutant subset (Supplementary Figure 11e). This relationship was consistently observed across multiple cancer types (Supplementary Figure 11f). Moreover, TP53-mutant tumors with high NuS exhibited significantly lower RiboSis than those with low NuS (Supplementary Figure 11g), further supporting a robust association between NuS and nucleolar functional state even in the absence of intact p53 signaling. These results indicate that although NuS is closely associated with p53-related signaling, it is not simply a surrogate for p53 activity, but reflects broader nucleolar functional states beyond the canonical p53 axis. Collectively, these analyses establish NuS and RiboSis as complementary and non-redundant metrics of nucleolar function. Their integration defines a dual-axis framework that captures both biogenesis activity and stress signaling, providing mechanistic insight into nucleolar regulation and prognostically informative stratification across human cancers.

### NuS-based screening identifies drugs inducing nucleolar stress

The mechanisms of action for many chemotherapeutics remain incompletely understood, limiting the interpretation of treatment efficacy, resistance, and toxicity. Accumulating evidence, including our findings with oxaliplatin, suggests that nucleolar stress can be an important but underappreciated component of drug response. This motivated a systematic effort to identify additional compounds capable of inducing nucleolar stress (Figure 7a).

**Figure 7.**
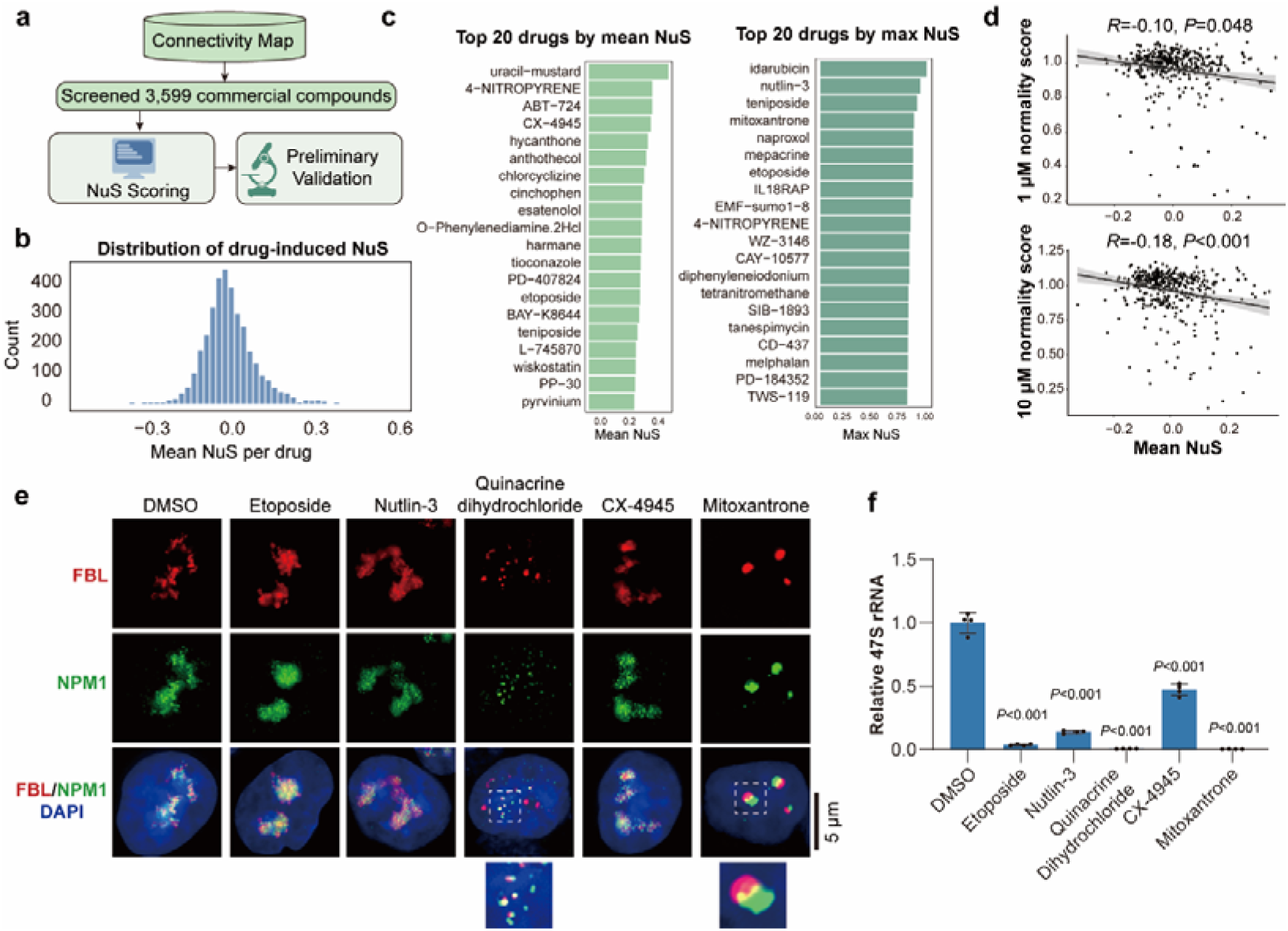
CMap-based identification of compounds associated with nucleolar stress. (a) Screening workflow for NuS-based analysis of CMap perturbational transcriptomes and experimental validation of prioritized compounds. (b) Distribution of mean NuS across all screened compounds. (c) Top 20 compounds ranked by mean NuS (left) or maximum NuS (right). (d) Correlation between mean NuS and morphological normality score at 1 μM and 10 μM. Each dot represents one compound; Spearman correlation coefficients and P values are shown. (e) SIM imaging of nucleolar morphology in HCT116 cells after treatment with the indicated compounds for 24 h. Cells were stained for FBL (red), NPM1 (green), and DAPI (blue). Insets show enlarged nucleoli. Scale bar, 5 μm. (f) RT-qPCR analysis of 47S pre-rRNA levels after compound treatment. Statistical significance was assessed by one-way ANOVA followed by Dunnett’s multiple comparisons test against the DMSO group; exact *P* values are shown. Source data for d and f are provided in Source Data file.

We analyzed Level 5 perturbational gene expression profiles for 3,599 compounds from the Connectivity Map (CMap) and computed NuS for each profile using GSVA (Supplementary Data 4). Drug-induced NuS values were approximately normally distributed (Figure 7b). To prioritize candidates, compounds were ranked based on both their mean NuS and maximum NuS across profiles, capturing both consistent and context-specific stress-inducing potential (Figure 7c).

Among the top-ranked compounds were multiple agents with established anti-tumor activity, as well as compounds used in anti-infective, neurological, and other therapeutic contexts. Candidate anti-cancer drugs represented diverse mechanistic classes, including DNA-alkylating agents (e.g., uracil-mustard and melphalan), topoisomerase II inhibitors (e.g., etoposide, teniposide, and mitoxantrone), anthracycline-related compounds (e.g., idarubicin and mitoxantrone), and targeted regulatory inhibitors such as CX-4945, Nutlin-3, and tanespimycin. Notably, the screen identified Nutlin-3 and etoposide, both previously reported to be associated with nucleolar stress responses ^41,42^, supporting the validity of the screening strategy. With the exception of topoisomerase inhibitors, for which links to nucleolar stress have been reported ^42–44^, the extent to which nucleolar stress contributes to the mechanisms of many other prioritized compounds remains unclear. This highlights the potential of NuS-based screening to uncover previously underappreciated aspects of drug action. To further evaluate the relationship between NuS-based screening and the previously established morphological normality score ^16^, we analyzed the drugs shared between the two datasets. Higher NuS was associated with lower normality scores, and this inverse correlation was more pronounced at 10 μM than at 1 μM, suggesting that the association strengthens under higher drug exposure (Figure 7d). Although the overall correlation was modest, these findings further support NuS as a reliable tool for the quantitative assessment of nucleolar stress.

To experimentally validate prioritized candidates, we selected representative compounds for testing in HCT116 cells, including Nutlin-3, etoposide, mitoxantrone, CX-4945, and quinacrine dihydrochloride (mepacrine salt). After 24 h of treatment, mitoxantrone and quinacrine dihydrochloride induced pronounced nucleolar morphological alterations, whereas the remaining compounds produced minimal structural changes under these conditions (Figure 7e). We reasoned that these compounds may induce weaker or slower nucleolar remodeling. Besides, RT-qPCR showed that all tested compounds significantly reduced 47S pre-rRNA levels (Figure 7f), indicating that nucleolar function was impaired and that inhibition of rRNA transcription can precede overt nucleolar structural disruption. This finding further underscores the sensitivity of NuS in detecting nucleolar stress. Together, these results demonstrate that NuS-based analysis of perturbational transcriptomes enables systematic mapping of nucleolar stress across diverse compounds, providing a framework for elucidating drug mechanisms and anticipating nucleolar-associated toxicities.

## Discussion

Quantifying nucleolar stress is essential for understanding its roles and mechanisms across disease contexts. Existing approaches largely rely on nucleolar morphology and a limited set of functional detection, including impaired nascent rRNA synthesis and processing, reduced protein synthesis, and activation of p53 signaling. Although informative, these approaches typically require dedicated experiments and are not easily scalable, limiting their applicability for systematic analyses across large cohorts and diverse data modalities. Here, we address this gap by curating nucleolar stress-associated gene sets and developing a quantitative nucleolar stress score (NuS) based on expression profiles. Across bulk transcriptomics, proteomics, single-cell, and spatial transcriptomic datasets, NuS robustly captured nucleolar stress signatures and complemented experimental assays, enabling more scalable and quantitative assessment of nucleolar stress across biological contexts.

Importantly, NuS should be interpreted as an associative metric of nucleolar stress, rather than a direct or causal measurement of nucleolar stress itself. Because it is derived from expression profiles, NuS reflects upstream regulatory inputs and downstream molecular responses associated with nucleolar perturbation, rather than directly measuring nucleolar disruption. Accordingly, changes in NuS are not expected to map in a strictly synchronous manner onto experimental indicators of nucleolar stress; depending on the cellular context, perturbation, and timing, they may precede, accompany, or follow detectable typical functional alterations such as impaired rRNA synthesis. An elevated NuS therefore suggests that cells may be undergoing, or have engaged programs consistent with, nucleolar stress, but it does not by itself constitute direct evidence of nucleolar dysfunction. This distinction is important because nucleolar stress is a dynamic and multifaceted process whose transcriptional changes vary across biological contexts.

Our analysis of 5-FU illustrates a key advantage of NuS. In HCT116 cells, NuS increased after 24 and 48 h of treatment, yet overt nucleolar morphological changes were not detectable at 24 h despite measurable suppression of rRNA transcription. This dissociation suggests that 5-FU can perturb rRNA synthesis without immediately triggering the structural remodeling typically used as a morphological hallmark of nucleolar stress, and that stronger or more prolonged inhibition may be required for morphological manifestation. This interpretation is further supported by our validation panel, in which Nutlin-3, etoposide, barasertib-HQPA, and CX-4945 markedly suppressed 47S pre-rRNA at 24 h but did not consistently induce obvious nucleolar structural changes. In these settings, NuS detected an early molecular stress response that precedes visible nucleolar disruption, highlighting its sensitivity. We also observed cell line-dependent responses: in HCT8 cells, 5-FU did not markedly inhibit rRNA transcription under our experimental conditions. This observation should be interpreted cautiously, as prior studies suggest that 5-FU may exert its nucleolar effects predominantly through disruption of rRNA processing rather than through strong inhibition of rRNA transcription.

A central implication of our work is that nucleolar stress and ribosome biogenesis are not simply inversely coupled. Reduced ribosome biogenesis activity is often interpreted as a substitute for elevated nucleolar stress; however, our analyses show that these two axes can vary independently. This motivates the combination of RiboSis and NuS as complementary metrics of nucleolar function. RiboSis reflects ribosome biogenesis capacity and biosynthetic demand, whereas NuS captures the stress response associated with maintaining nucleolar function under perturbation. In many settings, cells may sustain ribosome biogenesis to meet ongoing demands while concurrently experiencing elevated nucleolar stress, reflecting a balance between biosynthetic drive and stress burden. Notably, this does not imply that NuS directly measures the magnitude of nucleolar damage; rather, it indicates the extent to which nucleolar stress-related programs are engaged.

This duality is particularly relevant in cancer. Tumor cells frequently upregulate ribosome biogenesis to support rapid growth, yet they also experience chronic oncogenic, metabolic, and genotoxic pressures that can elevate nucleolar stress. Distinct nucleolar functional states defined by ribosome biogenesis activity and nucleolar stress may underlie therapeutic vulnerabilities in certain cancers. Indeed, classification of tumors using combined RiboSis and NuS revealed pronounced heterogeneity across cancer types and among patients within the same cancer type, consistent with contributions from both tumor-intrinsic programs and microenvironmental constraints. These nucleolar functional states were associated with patient outcomes in multiple cancers. While further validation in independent clinical cohorts is required, this framework may ultimately support prognosis prediction and guide therapeutic decision-making.

Spatial analyses further highlight the potential utility of expression-derived nucleolar metrics. In the CRC spatial datasets, RiboSis exhibited strong enrichment in tumor regions and delineated tumor niches with high specificity, suggesting practical value for spatial tumor mapping. Such a molecular signature could complement histopathology by providing an orthogonal, quantitative feature for tissue annotation and may be leveraged to support computational pathology workflows, for example by supplying molecularly informed labels for model training and benchmarking. In parallel, spatial variation in NuS may help identify regions with distinct nucleolar stress-related states, even when overt histological correlates are less apparent.

Finally, our NuS-based screen suggests that nucleolar stress is a broadly engaged drug response program and may be underappreciated for many compounds. Together with our observations in oxaliplatin-sensitive versus -resistant models, these findings support a model in which tumors with high ribosome biogenesis and relatively low baseline of nucleolar stress may be particularly susceptible to nucleolar stress-inducing therapies, whereas resistance may involve reduced ribosome biogenesis and/or blunted stress signaling. In this context, NuS provides a scalable approach to nominate candidate compounds that may exploit nucleolar vulnerabilities, and combined analysis of NuS and RiboSis may help prioritize drug mechanisms and therapeutic hypotheses for subsequent studies.

However, several limitations of this work should also be noted. First, NuS is a computational metric derived from expression profiles and therefore requires continued experimental validation across additional cellular contexts and perturbations. Second, score performance can be affected by incomplete gene coverage or platform-specific measurement biases, particularly in datasets with sparse profiles. Third, although the current Full and Core signatures capture key nucleolar stress biology, they may not fully encompass the complexity and context dependence of nucleolar regulation. Future studies integrating additional perturbational datasets, multi-omics evidence, and systematic genetic and pharmacological perturbations will further refine these signatures and improve the precision, interpretability, and biological scope of NuS and RiboSis as quantitative descriptors of nucleolar function.

## Methods

### Cell Culture

HCT8 and HCT116 cells were maintained in DMEM medium supplemented with 10% (v/v) fetal bovine serum (FBS) at 37°C in a humidified incubator with 5% CO_2_. Oxaliplatin-resistant derivatives of HCT116 and HCT8 (designated HCT116/L and HCT8/L, respectively) were generated by high-concentration pulse selection followed by stepwise dose escalation. Resistant lines were routinely maintained in medium containing oxaliplatin (2 μg/mL for HCT116/L and 1 μg/mL for HCT8/L). Before experiments, cells were cultured in drug-free medium for at least 1 week to minimize acute drug effects.

### Structured illumination microscopy (SIM) imaging

Cells were treated with the indicated compounds for 24 h and fixed in 4% paraformaldehyde (PFA) for 15 min at room temperature. After washing with ice-cold PBS, cells were permeabilized with 0.5% Triton X-100 for 5 min and blocked with 10% goat serum for 30 min. Samples were incubated overnight at 4 °C with primary antibodies against NPM1 (Cell Signaling Technology, #92825, 1:400) and fibrillarin (FBL; Abcam, #ab4566, 1:400). After PBS washes, fluorophore-conjugated secondary antibodies were applied for 1 h at 37 °C. Nuclei were counterstained with DAPI for 10 min at room temperature. Cells were mounted with a glycerol-based mounting medium and imaged on a Nikon structured illumination super-resolution microscope.

### Immunofluorescence (IF)

IF was used to examine the subcellular localization and abundance of specific proteins. HCT8 and HCT116 cells were treated with the indicated compounds and fixed with 4% paraformaldehyde (PFA) for 15 min at room temperature. After washing with ice-cold PBS, cells were permeabilized with 0.5% Triton X-100 for 5 min and blocked with 10% goat serum for 30 min. Samples were incubated overnight at 4 °C with primary antibodies against NPM1 (Cell Signaling Technology, #92825, 1:400). After PBS washes, fluorophore-conjugated secondary antibodies were applied for 1 h at 37 °C. Nuclei were counterstained with DAPI for 10 min at room temperature. Cells were mounted with a glycerol-based mounting medium and imaged on a confocal laser scanning microscope (Olympus FV4000). Fluorescence intensities were quantified in Fiji (ImageJ).

### RT-qPCR

Total RNA was extracted from cells after 24 h treatment with the indicated compounds using TRIzol reagent. cDNA was synthesized according to the manufacturer’s instructions using a reverse transcription kit (37°C for 15 min, 85°C for 5 s, then hold at 4°C). Dilute the cDNA 50 times with DEPC-treated water and used as template for SYBR Green-based qPCR with gene-specific primers. Primer sequences were as follows: 47S pre-rRNA, forward 5’-GAACGGTGGTGTGTCGTTC-3’ and reverse 5’-GCGTCTCGTCTCGTCTCACT-3’; GAPDH, forward 5’-GGAGCGAGATCCCTCCAAAAT-3’ and reverse 5’-GGCTGTTGTCATACTTCTCATGG-3’. Each 10 μL reaction contained 5 μL SYBR Green mix, 1 μL diluted cDNA, 0.4 μL forward primer (1 μM), 0.4 μL reverse primer (1 μM), and 3.2 μL DEPC-treated water. Reactions were run on a 7500 Fast Real-Time PCR System (Applied Biosystems). GAPDH served as the internal reference, and relative expression was calculated using the 2^-ΔΔCt^ method. Data are presented as mean ± s.d.

### Nascent RNA labeling with 5-ethynyluridine (5-EU) combined with IF

For nascent RNA labeling, HCT8 and HCT116 cells were incubated with 1 mM 5-ethynyluridine (5-EU; Beyotime Biotechnology, #R0309S-1) for 60 min. Cells were then fixed with 4% paraformaldehyde (PFA) for 15 min and permeabilized with 0.3% Triton X-100 for 15 min. If protein visualization was required, IF staining was performed as described above, including primary and secondary antibody incubations. After secondary antibody staining, EU was detected using a Click-iT reaction cocktail prepared according to the manufacturer’s instructions (86 μL Click Reaction Buffer, 4 μL CuSO₄, 0.2 μL Alexa Fluor 594 azide, and 10 μL Click Additive Solution) and incubated with samples for 30 min at room temperature in the dark. Samples were washed with PBS, nuclei were counterstained with DAPI for 10 min. Cells were mounted with a glycerol-based mounting medium and imaged on a confocal laser scanning microscope (Olympus FV4000). Fluorescence intensities were quantified in Fiji (ImageJ).

### 4D Fast DIA-based quantitative proteomics

HCT116 cells and the oxaliplatin-resistant derivative HCT116/L were treated with PBS or oxaliplatin for 24 h. Cells were washed twice with ice-cold PBS and lysed in four volumes of urea lysis buffer (8 M urea, 1% protease inhibitor cocktail, 3 μM trichostatin A, 50 mM nicotinamide). Lysates were sonicated on ice, clarified by centrifugation (12,000 × *g*, 10 min, 4°C), and protein concentrations were determined by BCA assay. Equal amounts of protein were precipitated with 20% TCA (final) at 4°C for 2 h, pelleted (4,500 × *g*, 5 min), washed 2-3 times with pre-chilled acetone, and air-dried. Pellets were resuspended in 200 mM triethylammonium bicarbonate (TEAB) and sonicated. Proteins were reduced with 5 mM DTT (56°C, 30 min), alkylated with 11 mM iodoacetamide (room temperature, 15 min, dark), and digested with trypsin (1:50, enzyme:protein) overnight at 37°C.

Peptides were dissolved in 0.1% formic acid (mobile phase A) and separated by nano-UHPLC (Vanquish Neo) using mobile phase B (0.1% formic acid in 80% acetonitrile) at 300 nL/min with a stepped gradient (4-22.5% B, 0-1.6 min; 22.5-35% B, 1.6-2.0 min; 35-55% B, 2.0-2.6 min; 55-99% B, 2.6-2.7 min; hold 99% B, 2.7-7.6 min). Eluted peptides were analyzed on an Orbitrap Astral mass spectrometer with nano-ESI (1.9 kV) in DIA mode. Full MS scans were acquired in the Orbitrap (m/z 480-780, 240,000 resolution), and MS/MS spectra were acquired in the Astral analyzer (starting m/z 150, 80,000 resolution) using sequential isolation windows and HCD (NCE 25). AGC target was set to 500% with a maximum injection time of 3 ms.

Raw DIA data were imported into database-search software and processed using analysis parameters defined according to the experimental design. DIA files were analyzed in Spectronaut (v18) using the Pulsar search engine with default settings. Searches were performed against the Homo_sapiens_9606_SP_20231220.fasta database (20,429 sequences) supplemented with a reverse decoy database to estimate false discovery rates (FDR). Up to two missed cleavages were allowed. The mass spectrometry proteomics data have been deposited to the ProteomeXchange Consortium (https://proteomecentral.proteomexchange.org) via the iProX partner repository ^45,46^ with the dataset identifier PXD072433.

### Data Analysis

#### Bulk Transcriptomic Datasets Analysis

##### Data collection and study design

Public bulk transcriptomic datasets were retrieved from the Gene Expression Omnibus (GEO) and selected to represent nucleolar stress induced by RNA polymerase I inhibition (CX-5461, BMH-21, or Act D). Only datasets with clearly annotated treatment conditions and matched controls were included. When multiple time points were available for the same cell line and compound, time points were analyzed jointly, and only genes that were differentially expressed at all time points were retained for downstream integration.

##### Data processing and gene annotation

Publicly available transcriptomic datasets were reanalysed according to the type of downloadable expression matrix provided in each study. For microarray datasets (GSE6400, GSE12666, GSE33417, GSE62593 and GSE62963), processed expression matrices were imported into R and probe identifiers were mapped to official gene symbols using platform-specific annotation resources. All Affymetrix datasets were processed and normalized by the authors. Prior to downstream analysis, the data distribution and expression scale were assessed for each dataset, and log_2_ transformation was applied when necessary. For RNA-seq datasets (GSE118565, GSE198178, GSE255898, GSE261563, GSE267499, GSE267501, GSE282212, GSE282214, and GSE298220), gene-level count tables were imported into R and annotated to official gene symbols using the corresponding platform annotation. Low-expression genes were removed prior to analysis, typically by requiring expression values >1 in at least three samples. Processed TPM and FPKM matrices were then transformed as log_2_(x + 1) for downstream analysis. For all datasets, duplicated gene symbols were collapsed by retaining the feature with the highest median expression across samples.

##### Differential expression analysis

Differential expression analysis was performed in R using limma. Briefly, a linear model was fitted to the expression matrix with treatment group as the main factor using “lmFit” and the specified design matrix. Pairwise contrasts were defined with “makeContrasts” and applied using “contrasts.fit”, followed by empirical Bayes moderation with “eBayes”. Differential expression statistics were extracted with “topTable” using Benjamini-Hochberg false discovery rate (FDR) correction. Genes were considered differentially expressed if they met |log_2_FC| > 1 and adjusted *P* value (or *P* value) < 0.05.

For the Agilent time-course dataset GSE62963, the downloaded matrix had already been normalized to the pretreatment baseline (0 h) and was therefore treated as a baseline-normalized log_2_ change matrix. In this case, each time point was analysed separately using an intercept-only limma model with empirical Bayes moderation to test whether the mean log_2_ change differed from zero. Stable responsive genes were defined by combining effect size, replicate consistency and statistical significance, using the following criteria: absolute mean log_2_ change > 1, standard deviation of log_2_ change ≤ 0.5 across replicates, concordant direction of change, and nominal *P* value < 0.05. For datasets generated from the same cell line with multiple time points, only genes with consistent directionality across time points were retained to reduce time-specific noise.

##### Cross-dataset integration for nucleolar stress gene screening

DEG lists from individual datasets were harmonized by gene symbol. To prioritize robust nucleolar stress-associated genes, we selected genes that were consistently upregulated or downregulated across independent datasets and retained those recurring above a predefined frequency threshold. Detailed information for each dataset, including platform type, preprocessing status, statistical strategy and DEG numbers, is provided in Supplementary Data 2.

##### NuS scoring in bulk transcriptomes

To quantify nucleolar stress in bulk expression profiles, the NuS score was computed using gene set-based enrichment of upregulated and downregulated components of the curated signature. For small cohorts or single-dataset validations, single-sample gene set enrichment (ssGSEA) was used for improved stability; for large-scale screening, GSVA was used. The NuS was calculated as:

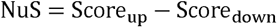

where Score_up_ and Score_down_ denote enrichment scores for the upregulated and downregulated gene subsets, respectively. A higher score indicates a higher level of nucleolar stress.

#### KEGG and Gene Ontology (GO) enrichment analysis

Functional enrichment analyses were performed to interpret the biological processes and pathways associated with the nucleolar stress gene sets. Gene Ontology (GO) (Biological Process, Molecular Function, and Cellular Component, as indicated) and Kyoto Encyclopedia of Genes and Genomes (KEGG) pathway enrichment were conducted in R using a hypergeometric over-representation test implemented in standard enrichment packages clusterProfiler. Gene identifiers were converted to Entrez IDs prior to KEGG analysis when required, and the background universe was defined as all genes detected in the corresponding expression datasets (or all protein-coding genes, where specified). Multiple-testing correction was performed using the Benjamini-Hochberg method, and terms with adjusted *P* < 0.05 were considered significant. Enrichment results were ranked by adjusted *P* value, and the top enriched terms were visualized using bar plots or dot plots).

#### DepMap gene dependency analysis

To assess the essentiality of nucleolar stress signature genes, gene dependency scores were obtained from the Cancer Dependency Map (DepMap) portal. CRISPR knockout-based gene effect scores were downloaded for all available cell lines and mapped to the genes in the Full and Core gene sets. For each gene, dependency scores across cell lines were aggregated and summarized to characterize overall essentiality. Genes were classified as broadly essential if their DepMap dependency scores were < −1 (more negative scores indicate stronger dependency). The proportion of essential genes was calculated for the Full and Core sets. Dependency score distributions for each gene set were visualized as density plots, with the < −1 region highlighted to facilitate comparison.

#### scRNA-seq analysis

Single-cell RNA-seq data form CRC were obtained from the Gene Expression Omnibus (GEO; GSE178341) and analyzed in R using Seurat (v4.3.0). We used the author-provided preprocessed Seurat expression matrix, for which quality control, doublet removal, and normalization had been performed in the original study. We then performed variable-feature selection, scaling, principal component analysis (PCA), shared nearest-neighbor graph construction, clustering, and uniform manifold approximation and projection (UMAP) based on the RNA assay. Major cell types and epithelial subpopulations were assigned according to the published annotation.

To quantify nucleolar stress and ribosome biogenesis at single-cell resolution, NuS and RiboSis were computed using Seurat ‘AddModuleScore’. For nucleolar stress, NuS were calculated separately for the upregulated and downregulated components of the nucleolar stress signature, and the final NuS was defined as: NuS = Score_up_ − Score_down_. RiboSis were computed using the published ribosome biogenesis gene set. These scores were compared across major cell lineages, epithelial subclusters, and tumor versus normal specimens. For epithelial cells, a dedicated subset object was generated and analyzed separately. Cell-cycle effects were evaluated using Seurat CellCycleScoring based on the updated 2019 S-phase and G2/M gene sets, and the influence of cell-cycle state was further assessed by phase-stratified analyses and regression of S.Score and G2M.Score.

#### Spatial RNA-seq analysis

Processed Spatial gene expression data of nine CRC specimens from four patients were obtained from http://biotrainee.cn:19195/CRLM-ST-share/. Downstream analyses were performed in R using Seurat (v4.3.0). Based on the H&E images provided by the authors and the corresponding clustering results, we re-clustered spots and performed manual re-annotation of spatial regions to refine subpopulation labels.

Gene set-based scoring was performed using Seurat ‘AddModuleScore’. NuS and RiboSis were calculated for each spot as described above and visualized on tissue sections to assess the spatial distribution of nucleolar stress and ribosome biogenesis across specimens.

#### TCGA data analysis

TCGA and GTEx RNA-seq data were obtained from the UCSC Xena platform using the Toil recompute TPM dataset together with matched phenotype and clinical annotations. Expression values were log_2_-transformed, Ensembl identifiers were mapped to gene symbols, and duplicate genes were collapsed by retaining the entry with the highest median expression. Samples were classified as TCGA tumor or GTEx normal, and only TCGA tumor and GTEx normal samples were retained for the primary analyses, with TCGA normal samples excluded. TCGA samples with evidence of prior treatment exposure were removed based on clinical annotations. Before scoring, genes with insufficient information were excluded, missing values were imputed using row medians, and genes with zero variance were removed. NuS and RiboSis were then calculated using ssGSEA, which was selected because it provides robust sample-level enrichment estimates and is well suited for pan-cancer analyses across heterogeneous tumor types. For NuS, enrichment scores for the upregulated and downregulated gene sets were first calculated separately. These scores were then standardized across all samples by subtracting the mean and dividing by the standard deviation, and the final NuS was defined as the standardized up-signature score minus the standardized down-signature score. RiboSis scores were standardized in the same manner to facilitate comparison across samples and downstream analyses. These normalized scores were subsequently used for pan-cancer tumor-versus-normal comparisons, subtype stratification, and survival analyses.

In CRC, CMS classification was performed using both CMScaller and CMSclassifier to enable cross-method comparison and assess the robustness of subtype-associated NuS and RiboSis patterns.b TP53 mutation status was obtained from the UCSC Xena TCGA Pan-Cancer (PANCAN) somatic mutation dataset, based on Mutation Annotation Format (MAF) files. Samples were classified as TP53-mutant if they harbored non-synonymous mutations in TP53, and as TP53 wild-type otherwise.

#### CMap data analysis

Level 5 gene expression profiles and compound metadata from the Expanded CMap LINCS Resource 2020 were obtained from ConnectivityMap (https://clue.io/data/CMap2020#LINCS2020). Compounds with an assigned chemical name or proprietary trade name were retained, yielding 3,599 compounds for downstream analysis. NuS were computed from Level 5 expression profiles using the GSVA R package (gsvaParam) with the curated NuS full gene set, and compounds were ranked by their mean and maximum NuS across profiles.

### Statistics and reproducibility

No statistical methods were used to predetermine sample size. Investigators were not blinded to group allocation during experiments or outcome assessment. Statistical analyses were performed in R (v4.3.0). For comparisons between two groups, either a two-sided Student’s t-test or a two-sided Wilcoxon rank-sum test was used, as appropriate. Comparisons among three or more groups were performed using one-way ANOVA followed by Tukey’s or Dunnett’s multiple-comparison test, as appropriate, or by the Kruskal-Wallis test for non-parametric data. Correlations were assessed using Spearman’s rank correlation coefficient. Survival differences were evaluated using the log-rank test, and multiple-testing correction was performed using the Benjamini-Hochberg method where indicated. For analyses with small sample sizes or non-normally distributed data, non-parametric methods were applied as appropriate. Exact statistical tests, sample sizes, and definitions of error bars are provided in the figure legends. *P* values are reported as exact values where possible.

### Data availability

The mass spectrometry-based proteomics dataset generated in this study has been deposited in the iProX repository under accession PXD072433. All public datasets used in this study are described in the relevant Methods sections, Supplementary Data, and corresponding figure legends. Source data are provided with this paper, and code used for the analyses is publicly available at https://github.com/JianxiongChen-cloud/nucleolar-stress-score.

## Acknowledgements

We thank the Dongguan People’s Hospital Research Startup Fund for supporting this project. This project is also supported by the National Natural Science Foundation of China (grant 82373066 to Jun Zhou and grant 82504182 to Jianxiong Chen) and the Guangdong Natural Science Foundation (Certificate No. 2026A1515011094 to Jianxiong Chen). We also thank the Morphology Platform of the Core Facility at Southern Medical University, in particular Ms. Manna Lin, Mr. Zhunqiang Zhong, Mr. Jianjun Li, Ms. Yanqiong Zeng, and Ms. Siqi Yang, for assistance with imaging.

## Author contributions

Conceptualization, Jianxiong Chen and Jun Zhou;

Methodology, Jianxiong Chen, and Jun Zhou;

Investigation, Jianxiong Chen, Shuai Xiao, Zhe Hao, Huimeng Xu, and Xuemei Xu;

Software, Jianxiong Chen, and Shuai Xiao;

Formal Analysis & Validation, Jianxiong Chen, Shuai Xiao, and Huimeng Xu;

Writing – Original Draft, Jianxiong Chen, and Zhe Hao;

Writing – Review & Editing, Jianxiong Chen, Zhe Hao, and Shuai Xiao;

Visualization, Jianxiong Chen;

Supervision, Jun Zhou;

Funding Acquisition, Jianxiong Chen and Jun Zhou.

