## Supplementary Information for "A nucleolar stress gene signature for quantitative scoring across multi-omics contexts"

**Supplementary Table 1.**

**Upregulated genes associated with nucleolar stress**

| **Gene Symbol** | **Other Name(s)** | **Reference(s)** |
| --- | --- | --- |
| AATF |  | 32587828 |
| AKT1 | AKT | 24525337 |
| ATF3 |  | 24098051 |
| ATM |  | 24675884, 37066425, 33146634, 27829214 |
| ATR |  | 40530693, 37066425, 33146634, 23447702, 25916852 |
| BAX |  | 36632465, 37645431 |
| BBC3 |  | 40343270 |
| CCDC137 |  | 40530693 |
| CDKN1A | p21 | 40639572, 39705402, 31061518 etc, wildy reported |
| CDKN1B | P27KIP1 | 28981686 |
| CDKN2A | p19ARF, p14ARF | 28820908, 39528457, 25691462, 14636574, 22467867 |
| CHEK1 | CHK1 | 23447702 |
| CNOT2 |  | 36569330 |
| CORO2B | coronin 2B | 38937439 |
| CTNNB1 | Beta-Catenin | 33221336 |
| E2F1 |  | 24675884 |
| EEF2 | eEF2 | 25332393 |
| EEF2K | eEF2K | 25332393 |
| EMG1 |  | 39333759 |
| FCN3 |  | 36632465 |
| GADD45A |  | 30177839 |
| HIF1A |  | 19680224, 17102617 |
| HNRNPK |  | 40338663 |
| MAP3K5 | ASK1 | 21383696 |
| MAP3K8 | TPL2 | 24998852 |
| MAPK14 | p38 | 21383696, 27445333 |
| MDM2 |  | 22777350, 38859834 |
| MDM4 | MDMX | 22777350 |
| NEAT1 |  | 28288210 |
| NCL |  | 16751805 |
| NOP53 | GLTSCR2 | 22522597 |
| NPM1 | NPM, B23 | 15144954 |
| KGD4 | MRPS36 | 17131359 |
| MYBBP1A |  | 21297583 |
| MYC | c-Myc | 30247545, 21383696, 21383696 |
| MYCN | N-Myc | 34262099 |
| NEDD8 |  | 22081073 |
| PAK1IP1 |  | 21097889 |
| PIM1 |  | 30247545 |
| PML |  | 15195100 |
| PRKAA1 | pAMPK | 39521100 |
| PRDM1 |  | 33972671 |
| RGS6 |  | 38409136 |
| RPL11 |  | 15152193, 12842086, 14612427, 16803902, 20194507, 21383696, 21807902, 21903592, 22262176, 22467867, 24141778, 26489471, 27975169, 30024791, 32198344, 22081073. 22588717 |
| RPL22 |  | 29207594, 30874462, 39116201, 39146182 |
| RPL23 |  | 31144295, 20554519, 29991511, 32236588, 15314174, 15314173, 22391559, 38981482 |
| RPL26 |  | 20542919, 18951086 |
| RPL3 |  | 25473889, 21705779, 23255119, 26636733, 32290083, 32244996, 28085118, 27835895, 27385096 |
| RPL4 |  | 26908445 |
| RPL5 |  | 22391559, 24141778, 32198344, 20554519, 35902361, 7935455, 22588717, 27265389 |
| RPL6 |  | 24174547 |
| RPS14 |  | 30706966 |
| RPS2 |  | 31928715, 32370049 |
| RPS25 |  | 22777350 |
| RPS27 |  | 21170087 |
| RPS27A |  | 21561866 |
| RPS27L |  | 21170087 |
| RPS7 |  | 30177839, 30247545, 19683495, 20554519 |
| RPS9 |  | 36071402 |
| SIRT7 |  | 32404984 |
| SNORA13 |  | 38981482 |
| SOCS3 |  | 23686487 |
| SRSF1 |  | 23478443, 24807918 |
| TOPBP1 |  | 25916852 |
| TP53 |  | 40726293, 40639572, 39705402 etc, wildly reported |
| WNT4 |  | 32817586 |

*References are listed with PubMed IDs (PMID).*

**Supplementary Table 2.**

**Downregulated genes associated with nucleolar stress**

| **Gene Symbol** | **Other Name(s)** | **Reference(s)** |
| --- | --- | --- |
| AGTPBP1 |  | 30905767 |
| AKT1 | AKT | 23459592, 26993775 |
| AKT1S1 | PRAS40 | 24704832 |
| BCL2 |  | 36632465, 37645431 |
| BOP1 |  | 33510838, 38409361 |
| BRIX1 |  | 39475053 |
| CCDC137 |  | 40530693 |
| CCND1 | Cyclin D1 | 31659203 |
| CDK4 |  | 30706966 |
| CDK6 |  | 30706966 |
| DHODH |  | 32034120 |
| DHX16 |  | 39333759 |
| DHX33 |  | 36631557 |
| E2F1 |  | 32535367, 33298840, 31659203, 35565259 |
| EGR1 |  | 24098051 |
| EIF2AK4 | GCN2 | 37452637 |
| EXOSC8 |  | 36348012 |
| FANCA |  | 33523834 |
| FBXO7 |  | 31144295 |
| FTO |  | 40623539, 40213545 |
| GNL3 | NS, Nucleostemin | 19033382, 21444791 |
| GRK5 |  | 33507833 |
| GRWD1 |  | 29991511, 28722511, 27856536 |
| HEATR1 |  | 31190896, 37247644, 29143558 |
| IGF1 |  | 20392698 |
| IMPDH2 |  | 25347121, 39235732, 31371825 |
| KMT5A | SETD8 | 39341827 |
| LAS1L |  | 34319761 |
| LIN28A |  | 34331666 |
| MDM2 |  | 23807770，38285632, 22051195, 33298840 |
| MDM4 |  | 17327702, 17110929, 38285632 |
| MRPL44 |  | 40623539 |
| MRPS16 |  | 40623539 |
| MTREX | MTR4 | 36403484 |
| MYC | c-myc | 20194507, 24141778, 39341827, 38169774, 28288210, 21807902, 21807902, 37247644, 36569330, 31061518 |
| NBN | NBS1 | 40067889 |
| NCL |  | 38409136 |
| NDC80 |  | 39705402 |
| NEDD8 |  | 25867069, 26993774 |
| NFE2L2 | NRF2 | 27445333 |
| NIFK |  | 25826659 |
| NOD2 |  | 24098051 |
| NOL12 |  | 30988155 |
| NOL7 |  | 37246770 |
| NOLC1 | NOPP140 | 38062753, 23412656, 34201772, 30063880 |
| NOP2 |  | 34319761 |
| NOP53 | PICT1 | 21804542, 16043506, 22320853, 36403484, 28722511, 27829214, 25464032, 24923447 |
| NPM1 |  | 16855788, 22166220, 30814495, 21444791, 14636574, 33221336 |
| NUF2 |  | 39705402 |
| NUMA1 |  | 28981686 |
| PELP1 |  | 34319761 |
| PIM1 |  | 20639905, 26993775 |
| POLR1A | RPA194 | 34572872, 31285544, 25344835 |
| POLR1G | PAF49 | 37356716 |
| PPAN |  | 25759387, 30716409, 33221336, 36762786 |
| PSMD9 |  | 34077860 |
| PYGO2 |  | 23517060 |
| RAP1GDS1 | SmgGDS | 28806394 |
| RBM28 |  | 34953860 |
| RIOK1 |  | 23459592, 29712692 |
| RIOK2 |  | 23459592 |
| RPL11 |  | 19129914, 27734913, 20221446 |
| RPL13 |  | 35137207 |
| RPL18 |  | 35137207 |
| RPL23 |  | 24061479, 22588717 |
| RPL27A |  | 21674502, 32535367 |
| RPL32 |  | 32516735 |
| RPL37 |  | 20935493 |
| RPL9 |  | 40623539, 40213545 |
| RPS14 |  | 22391559 |
| RPS15A |  | 40623539, 40239541 |
| RPS19 |  | 22391559, 23412928, 18515656, 27734913, 34757171 |
| RPS2 |  | 33069875 |
| RPS26 |  | 23728348 |
| RPS27A |  | 35073964 |
| RPS27L |  | 25144937 |
| RPS6 |  | 19287375, 27734913 |
| RPS6KA1 | RSKS-1 | 29567958 |
| RPS7 |  | 35871033, 19683495, 22588717 |
| RPS8 |  | 35137207 |
| RPS9 |  | 20221446 |
| RRP12 |  | 26499779 |
| RRP15 |  | 34343634, 28099941 |
| RRP8 |  | 29567958 |
| RRS1 |  | 34433556 |
| SBDS |  | 32198344, 36632465 |
| SCD |  | 36675264 |
| SIRT1 |  | 34343634, 30623565 |
| SPEN |  | 37607001 |
| SURF2 |  | 39333141 |
| TAF1B |  | 38169774, 37645431 |
| TCOF1 |  | 15249688 |
| TERF2 | TRF2 | 29725012 |
| TRIM24 | TIF1A | 24273493, 23764776, 37626880, 34572268, 36229381, 28360124, 29873780, 28334682 |
| UBTF | UBF | 30814495, 28806394, 26317157 |
| USP47 |  | 32370049 |
| UTP11 |  | 37087976 |
| UTP14A |  | 21078665 |
| UTP18 |  | 20056613 |
| WDR3 |  | 20392698 |
| WDR5 |  | 30865883 |
| WDR75 |  | 34611297 |
| YBX1 |  | 36632465 |

*References are listed with PubMed IDs (PMID).*

**Supplementary Table 3.**

**Proteins with altered subcellular localization in nucleolar stress response**

| **Gene Symbol** | **Other Name(s)** | **Reference(s)** |
| --- | --- | --- |
| CDKN2A | p14ARF | 39690184, 36230979, 16855788 |
| E2F1 |  | 24675884 |
| EIF6 |  | 36632465 |
| GNL3 | NS | 36230979, 21464199, 39690184 |
| HEY1 |  | 27129302 |
| MAPT | Tau | 35170061, 30064522 |
| MYBBP1A |  | 23583237, 21297583, 24710530, 24375404 |
| NCL |  | 21464199, 39690184 |
| NOLC1 |  | 38617541 |
| NPM1 |  | 24998852, 39690184, 38617541, 36825461, 36230979, 30301139 |
| PARP1 |  | 38861757 |
| PICT1 |  | 39690184 |
| PPAN |  | 25759387, 36762786 |
| RAD9B |  | 22399810 |
| RBM28 |  | 34953860 |
| REL1 | p65 | 36230979, 30301139, 27445333, 37247644 |
| RPL3 |  | 38617541 |
| RPL5 |  | 25464032, 17327702, 17242401, 32516735, 17329973, 39690184 |
| RPL6 |  | 24174547 |
| RPL11 |  | 25464032, 17327702, 17242401, 14612427, 30024791, 32516735, 17329973, 39690184, 38409361 |
| RPL23 |  | 17327702, 17242401M, 17329973, 19033382, 39690184 |
| RPS7 |  | 39690184 |
| RRN3 |  | 28334682 |
| RSL1D1 | CSIG | 27811966 |
| TERF2 | TRF2 | 38062753 |
| VHL |  | 36230979 |
| WDR12 | PeBoW | 27440937 |

*References are listed with PubMed IDs (PMID).*

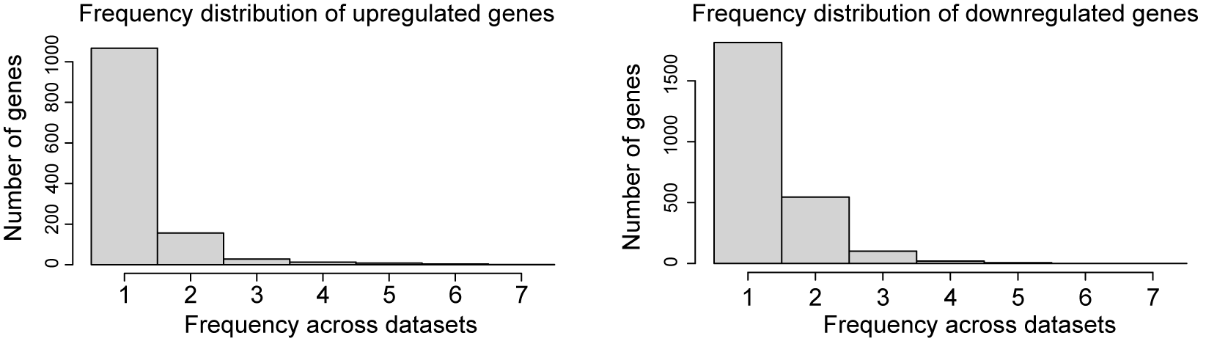
**Supplementary Figure 1.**

**Supplementary Figure 1. Frequency of differentially expressed genes (DEGs) across nucleolar stress datasets.** Histogram showing the distribution of DEG recurrence across 17 transcriptomic datasets, summarizing the occurrence frequencies of 1,277 upregulated and 2,489 downregulated genes.

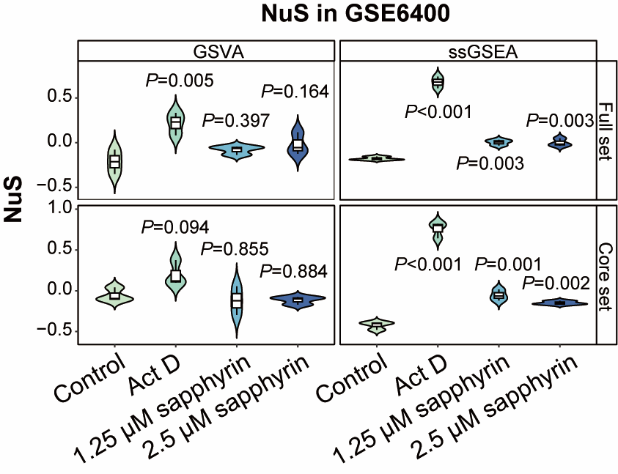
**Supplementary Figure 2.**

**Supplementary Figure 2. ssGSEA- and GSVA-based quantification of nucleolar stress in A549 cells.** NuS scores were calculated from GSE6400 transcriptomic data for A549 cells under the indicated treatments using ssGSEA and GSVA. Statistical analysis was performed by one-way ANOVA with Dunnett’s multiple-comparison test, using the control group as the comparator for all treatment groups. Exact *P* values are indicated in the figure.

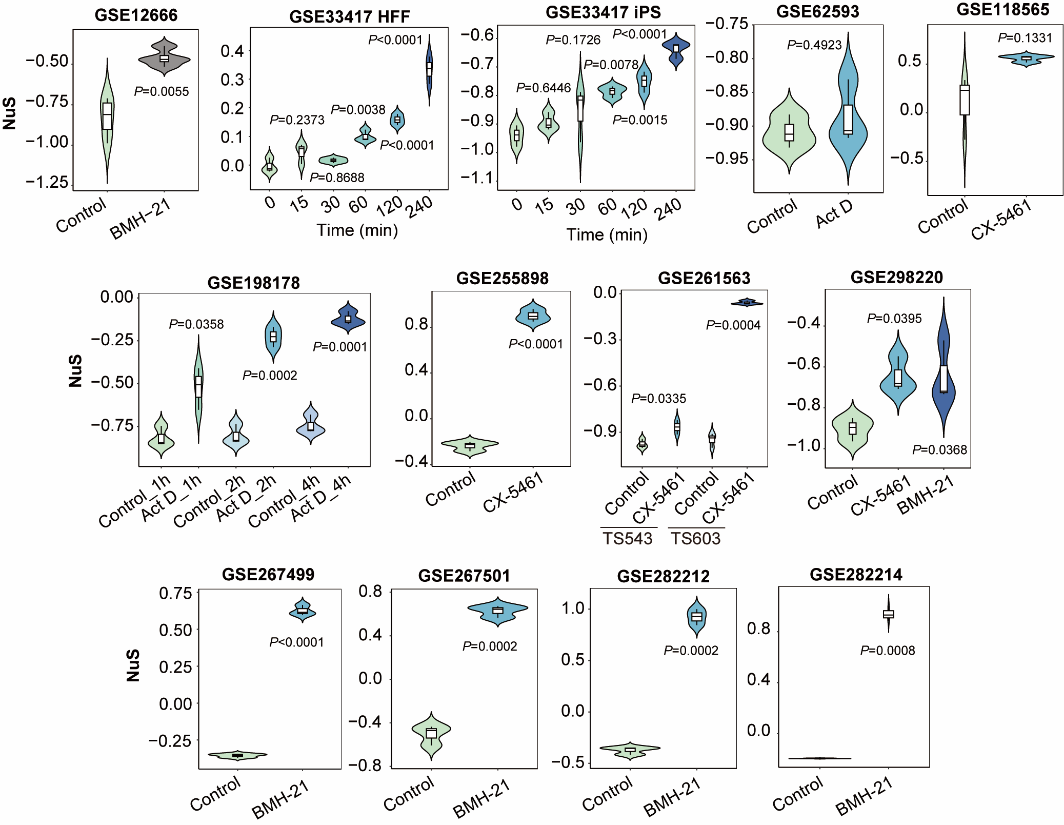
**Supplementary Figure 3.**

**Supplementary Figure 3. Internal validation of the NuS across nucleolar stress datasets.** NuS scores were calculated by ssGSEA with Full Set across 13 transcriptomic datasets. Statistical analysis was performed using two-sided Welch’s t-test for two-group comparisons, or one-way ANOVA with Dunnett’s multiple-comparison test for multi-group comparisons, using untreated control samples as the reference. Exact *P* values are shown.

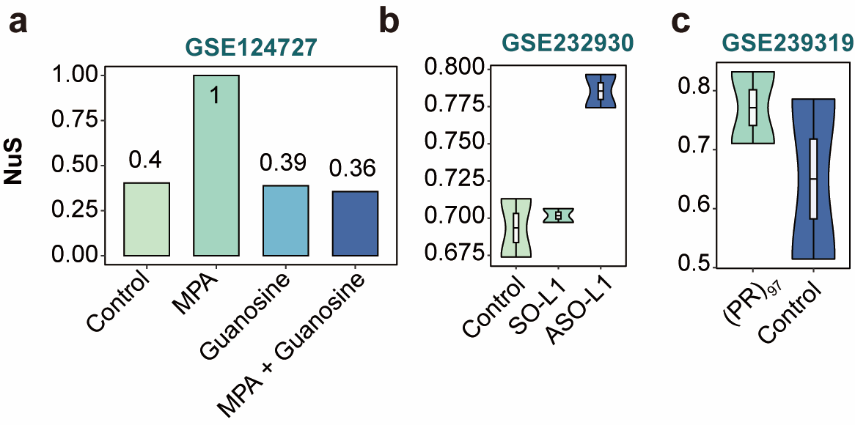
**Supplementary Figure 4.**

**Supplementary Figure 4. NuS across independent public datasets.** (a) GSE124727. NuS scores in cells treated with mycophenolic acid (MPA), guanosine, or MPA plus guanosine. (b) GSE232930. NuS scores in cells treated with a sense oligonucleotide (SO-L1) or a LINE1-targeting antisense oligonucleotide (ASO-L1). (c) GSE239319. NuS scores in samples with inducible expression of (PR)^97^ and corresponding controls. Owing to the limited sample size (*n* = 1 in a; *n* = 2 per group in b and c), no statistical analysis was performed; the plots are shown for descriptive comparison only.

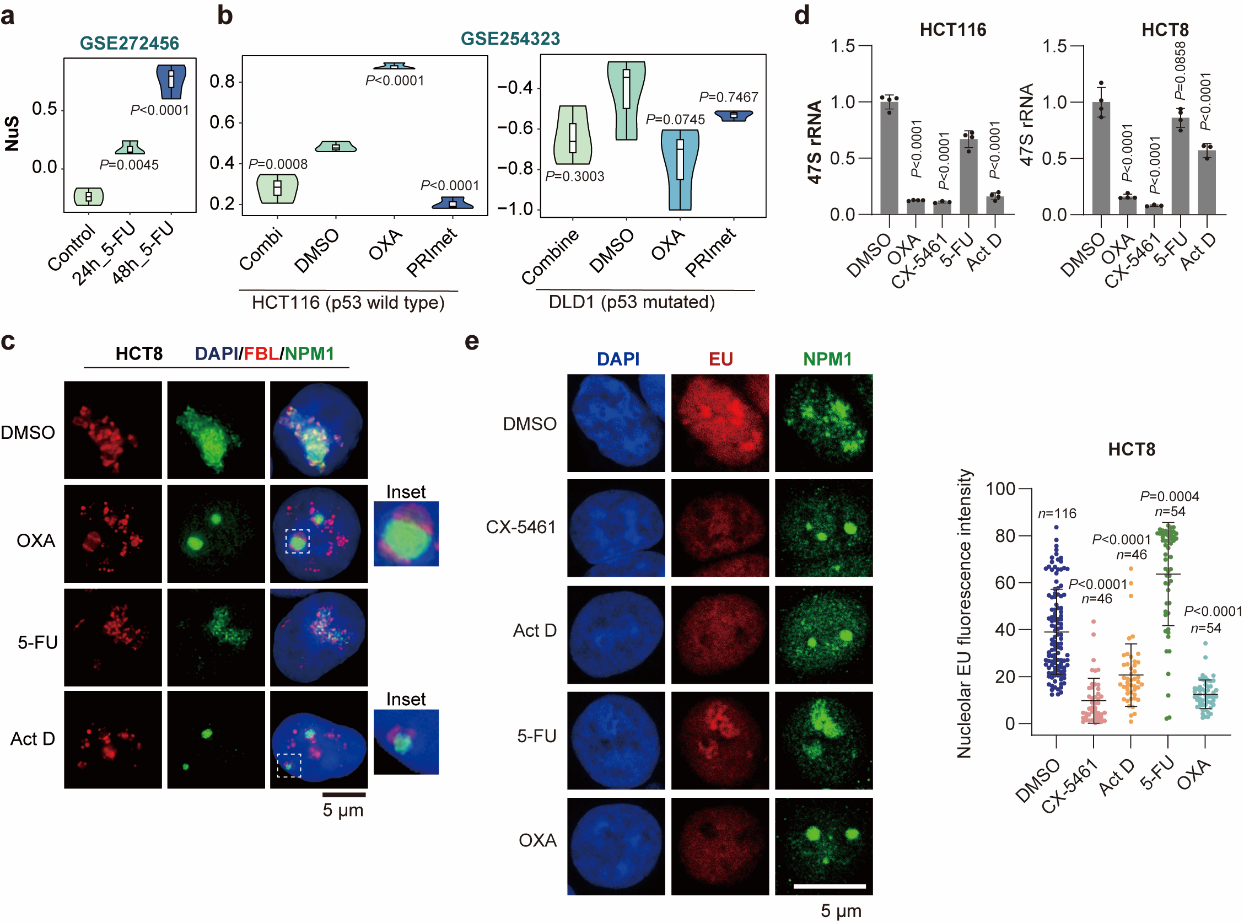
**Supplementary Figure 5.**

**Supplementary Figure 5. Chemotherapeutic agents increase NuS and suppress rRNA transcription.** (a) GSE272456. NuS scores in cells treated with 5-FU for 24 h or 48 h. Statistical analysis was performed using one-way ANOVA followed by Dunnett’s multiple-comparison test, with the control group as the reference. (b) GSE254323. NuS scores in DLD1 (p53-mutant) and HCT116 (p53-wild-type) cells treated with oxaliplatin (OXA) or PRImet. Statistical analysis was performed using one-way ANOVA followed by Dunnett’s multiple-comparison test, with DMSO as the reference group for each cell line. (c) SIM imaging of fibrillarin (FBL, red) and NPM1 (green) in HCT8 cells treated with DMSO, OXA, 5-FU or Act D. (d) qPCR analysis of 47S pre-rRNA levels in HCT116 and HCT8 cells treated with OXA, CX-5461, 5-FU or Act D. Statistical analysis was performed using one-way ANOVA followed by Dunnett’s multiple-comparison test, with DMSO as the reference group. (e) 5-EU incorporation assay in HCT8 cells treated with the indicated agents. Representative images show 5-EU (red) and NPM1 (green). Right, quantification of nucleolar EU fluorescence intensity; each dot represents one nucleolus, with n indicated. Statistical significance was assessed using the Kruskal–Wallis test. Exact *P* values are shown in the figure. Scale bar, 5 μm. Source data for d and e are provided in Source Data file.

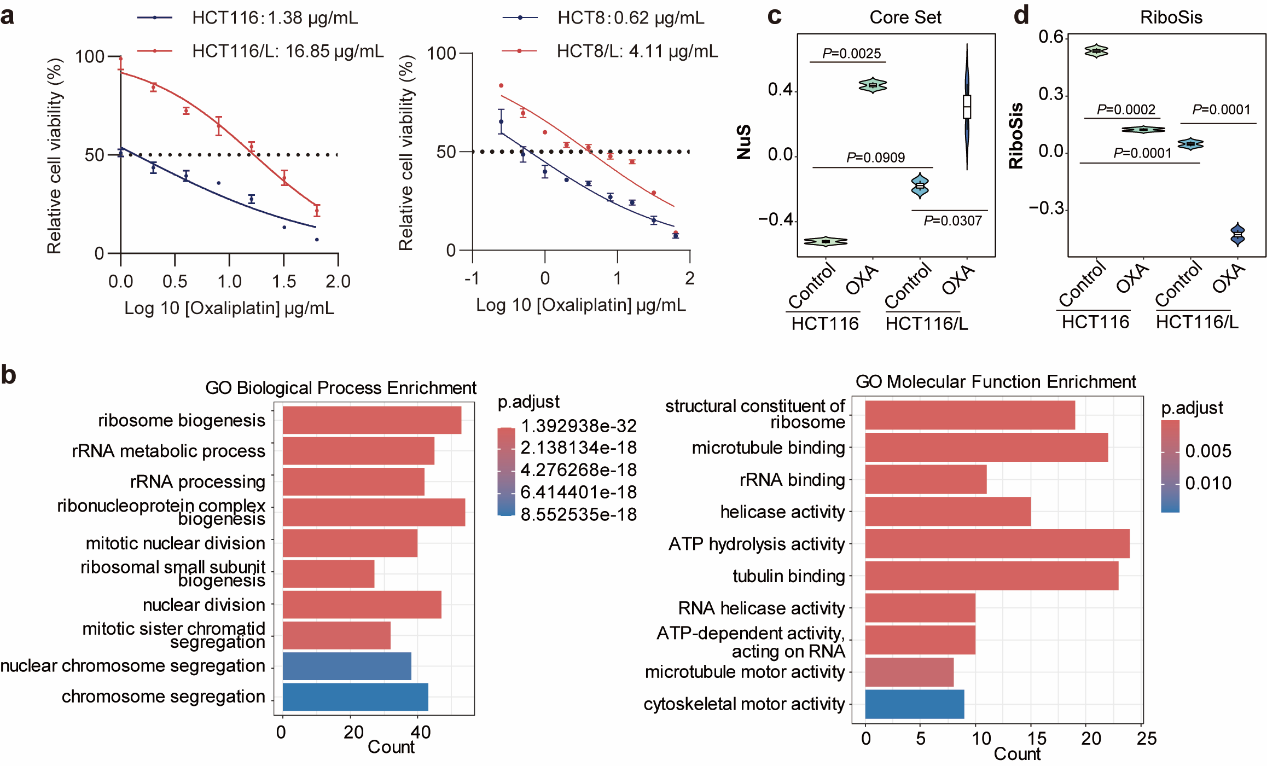
**Supplementary Figure 6.**

**Supplementary Figure 6. Oxaliplatin sensitivity, pathway enrichment, and RiboSis changes in parental and resistant CRC cells.** (a) Dose-response curves showing relative viability of HCT116, HCT116/L (oxaliplatin-resistant), HCT8, and HCT8/L cells following oxaliplatin treatment. (b) GO enrichment analysis of differentially expressed genes in oxaliplatin-treated HCT116 cells. (c–d) ssGSEA-based NuS (Core Set) and RiboSis scores in parental (HCT116) and resistant (HCT116/L) cells under control and oxaliplatin (OXA) treatment. Data are shown as violin plots with embedded boxplots. Statistical analysis was performed using one-way ANOVA followed by Tukey’s multiple comparisons test for multi-group comparisons. Exact *P* values are indicated in the figure. Source data for a, c, and d are provided in Source Data file.

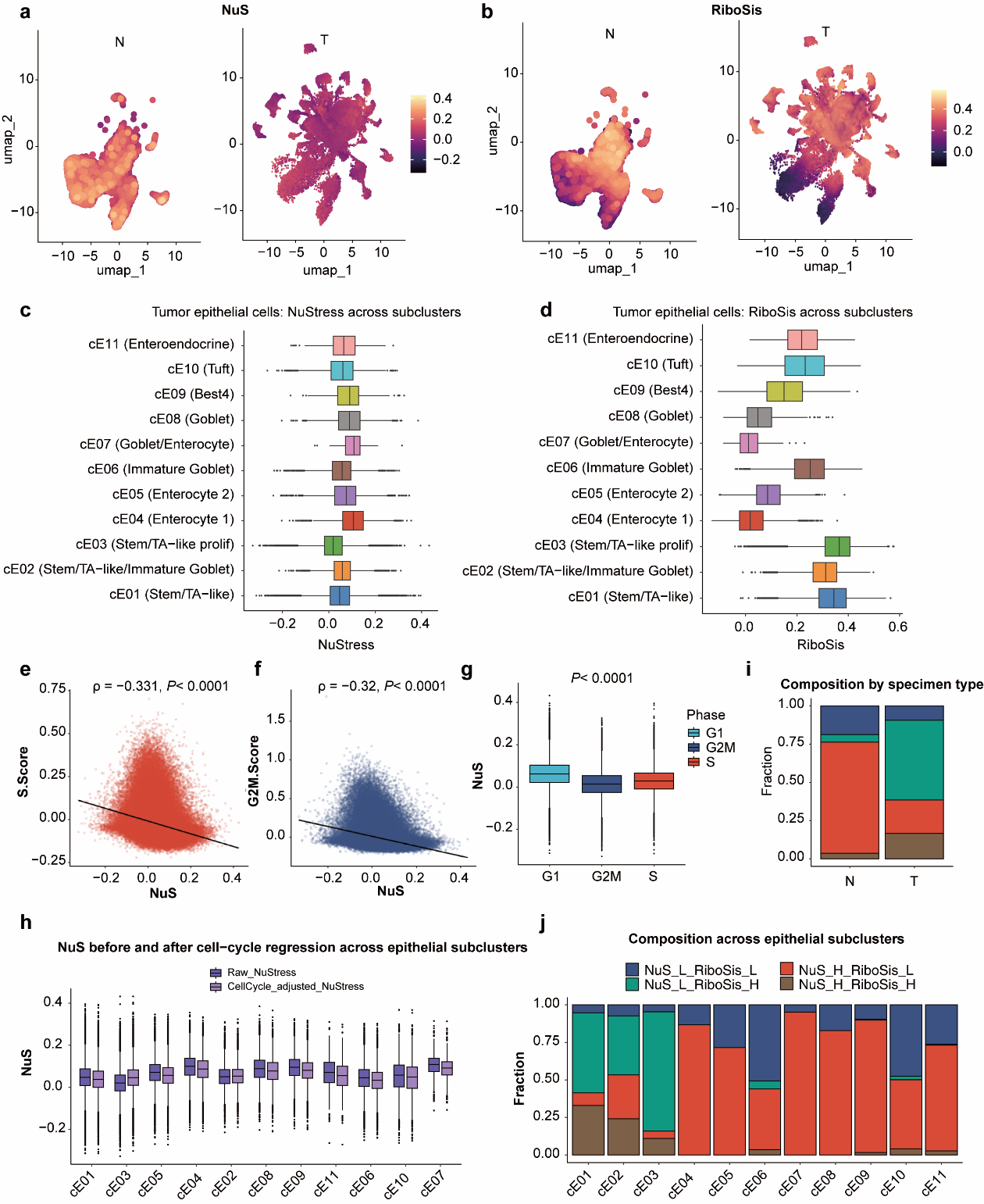
**Supplementary Figure 7.**

**Supplementary Figure 7. Cell-cycle effects and epithelial heterogeneity of NuS and RiboSis at single-cell resolution.** (a–d) Distribution of NuS (a,c) and RiboSis (b,d) across epithelial subpopulations. UMAPs show the spatial distribution of NuS (a) and RiboSis (b) in normal (N) and tumor (T) epithelial cells, and box plots summarize NuS (c) and RiboSis (d) across epithelial subclusters. (e,f) Spearman correlation analysis between NuS and cell-cycle scores. NuS showed a modest negative correlation with S-phase score (e) and G2/M score (f). Spearman’s rank correlation coefficient (ρ) and corresponding *P* values are indicated. (g) Comparison of NuS across cell-cycle phases (G1, S, and G2/M). Statistical significance was assessed using the Kruskal–Wallis test. (h) Box plots showing NuS before and after regression of cell-cycle effects across epithelial subpopulations. Adjusted NuS was obtained by regressing out S.Score and G2M.Score and re-centering the residuals, indicating that epithelial subcluster-specific variation in NuS remained evident after cell-cycle correction. (i,j) Composition of NuS/RiboSis-defined groups in normal and tumor epithelial cells (i) and across epithelial subpopulations (j).

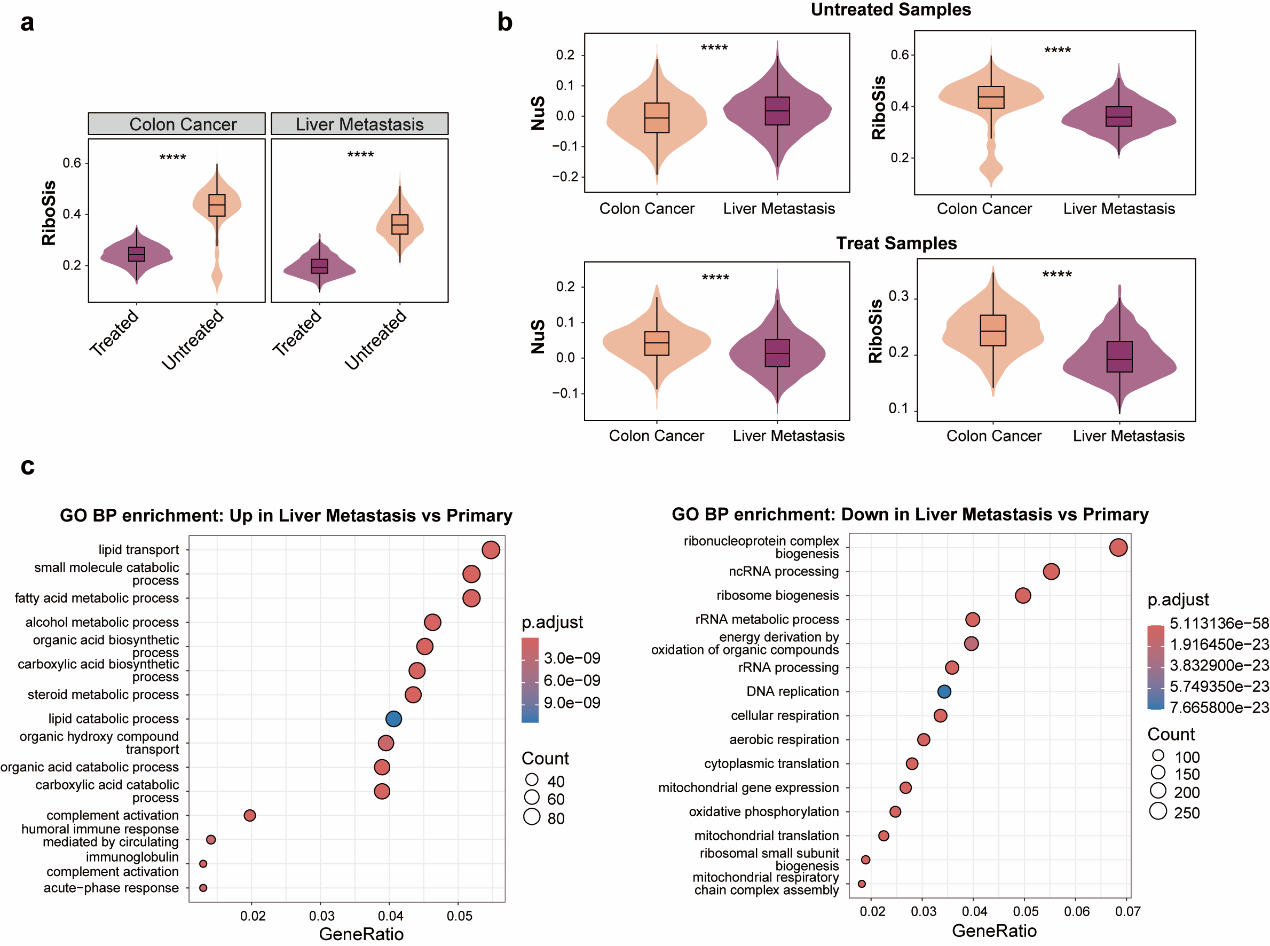
**Supplementary Figure 8.**

**Supplementary Figure 8. NuS and RiboSis in primary tumors and liver metastases with and without neoadjuvant treatment.** (a) Violin plots with overlaid box plots showing RiboSis in tumor regions from untreated and neoadjuvant-treated samples in primary tumors and matched liver metastases. Statistical significance was assessed using two-sided Wilcoxon rank-sum tests. (b) Violin plots with overlaid box plots comparing NuS between primary tumors and liver metastases within untreated and neoadjuvant-treated groups. Statistical significance was assessed using two-sided Wilcoxon rank-sum tests. (c) Gene Ontology (GO) biological process enrichment analysis of differentially expressed genes between liver metastases and primary tumors. Genes upregulated in liver metastases (left) and downregulated genes (right) are analyzed. Differential expression analysis was performed using Seurat FindMarkers with a two-sided Wilcoxon rank-sum test, and GO enrichment was conducted using clusterProfiler with Benjamini-Hochberg correction. **** *P*<0.0001.

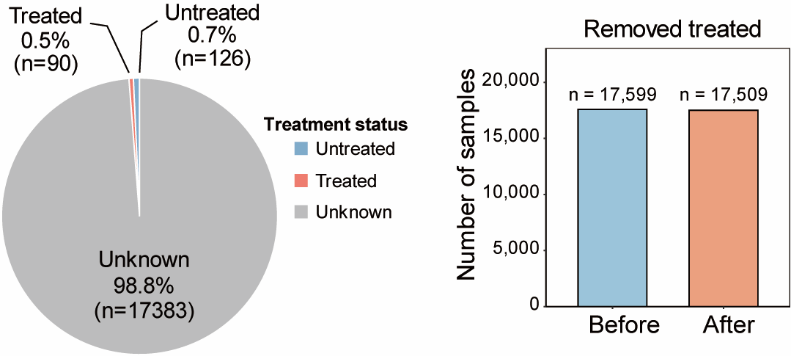
**Supplementary Figure 9.**

**Supplementary Figure 9. Filtering of treatment-exposed samples.** Proportion of untreated, treated, and treatment-unknown samples before filtering (left). Sample numbers before and after exclusion of treatment-exposed samples (right). Samples annotated as treated were removed to minimize confounding in downstream analyses.

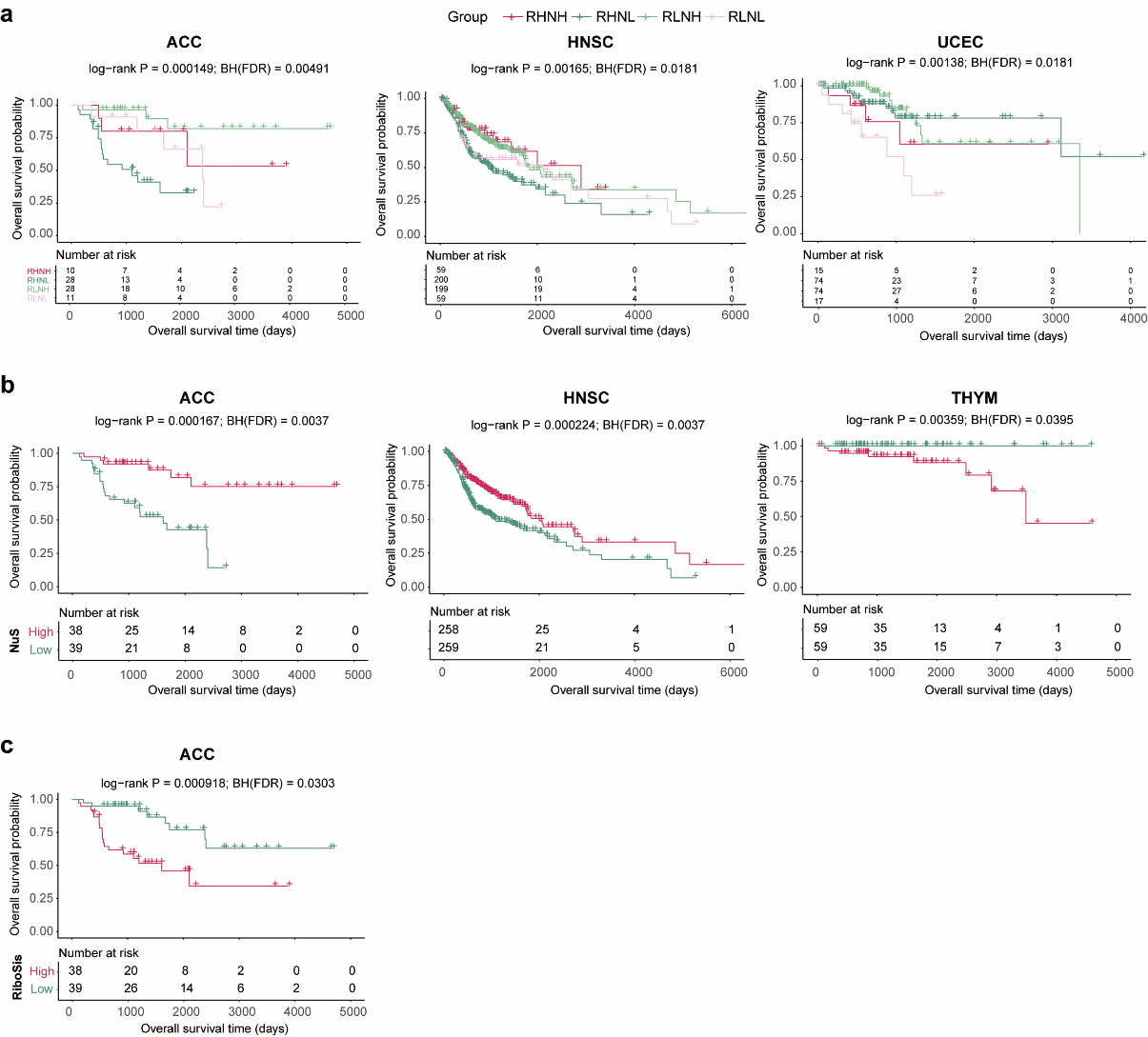
**Supplementary Figure 10.**

**Supplementary Figure 10. Prognostic value of NuS and RiboSis in overall survival analysis.** (a) Kaplan–Meier overall survival curves for patients stratified by the combined nucleolar functional states defined by NuS and RiboSis. Patients were classified into four groups using median cutoffs of NuS and RiboSis. (b) Kaplan–Meier overall survival curves for patients stratified by NuS alone (high vs low, median cutoff). (c) Kaplan–Meier overall survival curves for patients stratified by RiboSis alone (high vs low, median cutoff). Overall survival differences between groups were assessed using the log-rank test. Corresponding P values and Benjamini–Hochberg (BH)-adjusted P values are indicated in the plots. Hazard ratios (HRs) and 95% confidence intervals (CIs) were estimated using Cox proportional hazards models.

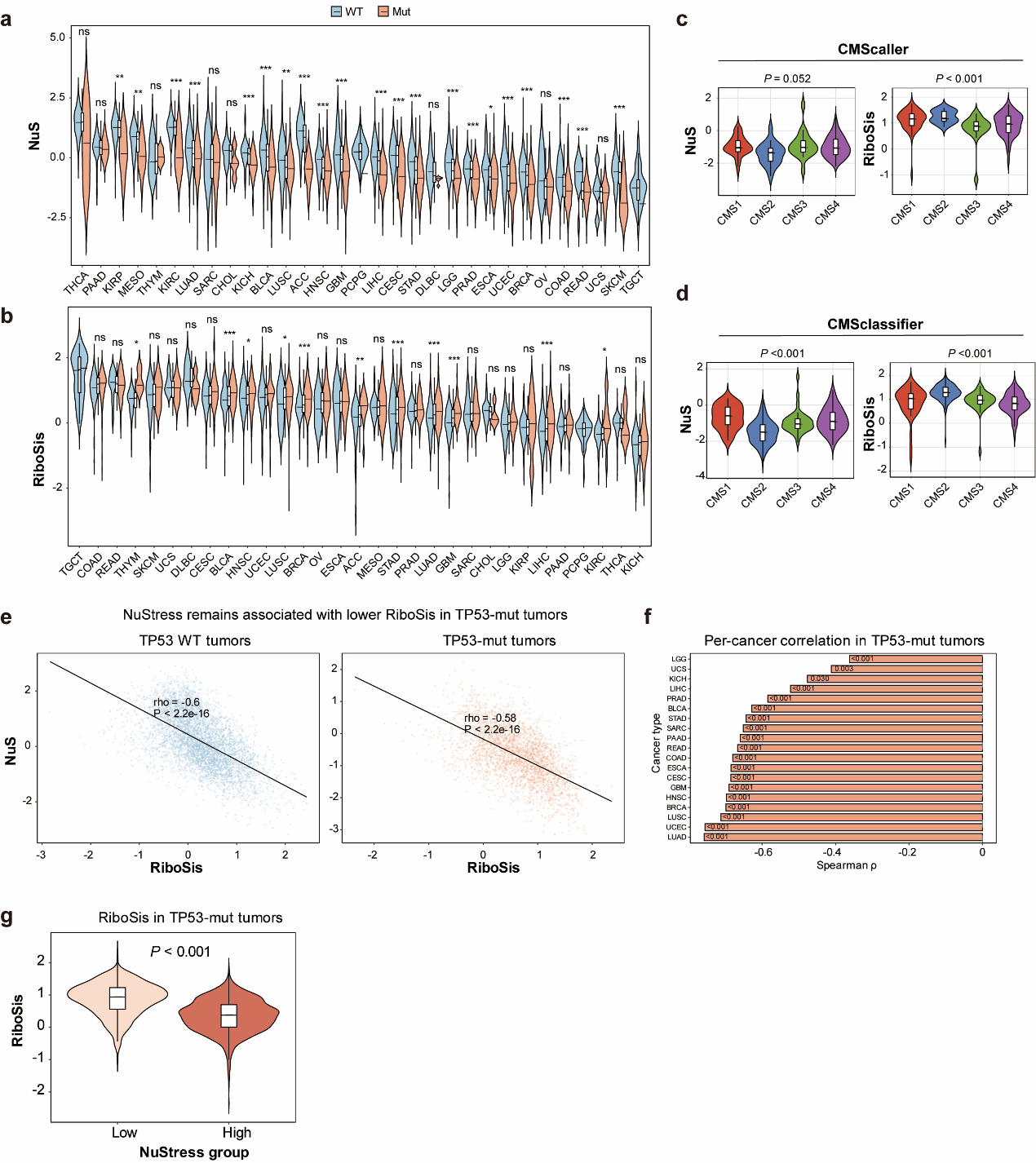
**Supplementary Figure 11.**

**Supplementary Figure 11. Association of NuS and RiboSis with TP53 status and CRC subtypes.** (a, b) Comparison of NuS (a) and RiboSis (b) between TP53 wild-type (WT) and TP53-mutant (Mut) tumor samples across cancer types. Statistical significance was assessed using a two-sided Wilcoxon rank-sum test. (c, d) Association of NuS and RiboSis with colorectal cancer (CRC) consensus molecular subtypes (CMS) defined using two classification methods, CMScaller (c) and CMSclassifier (d). NuS and RiboSis were compared across CMS groups. Statistical significance was evaluated using the Kruskal–Wallis test followed by post hoc multiple-comparison testing. (e) Correlation between NuS and RiboSis in TP53 wild-type and TP53-mutant tumor samples. Each dot represents one sample; Spearman correlation coefficients (ρ) and corresponding P values are shown. (f) Per-cancer correlation analysis between NuS and RiboSis in TP53-mutant tumors. Bars indicate Spearman correlation coefficients for individual cancer types. (g) Comparison of RiboSis levels between NuS-defined groups (high versus low) in TP53-mutant tumors. Statistical significance was assessed using a two-sided Wilcoxon rank-sum test. ns, not significant; * *P* < 0.05; ** *P* < 0.01; *** *P* < 0.001.

**Supplementary Data 1. Literature-curated nucleolar stress-responsive genes.**

Literature-curated nucleolar stress-responsive genes, including gene symbol, direction of change, and supporting reference(s).

**Supplementary Data 2. Differential expression results for 17 transcriptomic datasets.**

Differential expression results from 17 transcriptomic datasets treated with RNA polymerase I inhibitors, including log_2_ fold change, *P* value, adjusted *P* value (where available), and direction of regulation.

**Supplementary Data 3. Full list of differentially expressed genes identified across all 17 datasets.**

Combined list of differentially expressed genes identified across the 17 datasets before frequency-based filtering.

**Supplementary Data 4. Dataset-derived nucleolar stress gene set after frequency filtering.**

Dataset-derived nucleolar stress-associated genes recurrently altered in at least four independent datasets.

**Supplementary Data 5. Full Set of nucleolar stress-associated genes.**

Integrated Full Set of nucleolar stress-associated genes derived from literature curation and dataset-based screening after removal of discordant genes.

**Supplementary Data 6. Core Set of high-confidence nucleolar stress-associated genes.**

High-confidence Core Set of nucleolar stress-associated genes defined using more stringent literature- and dataset-based criteria.
